# Seabird and sea duck mortalities were lower during the second breeding season in eastern Canada following the introduction of Highly Pathogenic Avian Influenza A H5Nx viruses

**DOI:** 10.1101/2024.05.03.591923

**Authors:** Tabatha L. Cormier, Tatsiana Barychka, Matthieu Beaumont, Tori V. Burt, Matthew D English, Jolene A. Giacinti, Jean-François Giroux, Magella Guillemette, Kathryn E. Hargan, Megan Jones, Stéphane Lair, Andrew S. Lang, Christine Lepage, William A. Montevecchi, Ishraq Rahman, Jean-François Rail, Gregory J. Robertson, Robert A. Ronconi, Yannick Seyer, Liam U. Taylor, Christopher R. E. Ward, Jordan Wight, Sabina I. Wilhelm, Stephanie Avery-Gomm

**Affiliations:** Environment and Climate Change Canada, Canadian Wildlife Service, Sackville, New Brunswick, Canada; Environment and Climate Change Canada, Wildlife and Landscape Science Directorate, Ottawa, Ontario, Canada; Environment and Climate Change Canada, Canadian Wildlife Service, Québec, Quebec, Canada; Memorial University of Newfoundland, Department of Psychology, St. John’s, Newfoundland and Labrador, Canada; Environment and Climate Change Canada, Canadian Wildlife Service, Dartmouth, Nova Scotia, Canada; Université du Québec à Montréal, Département des Sciences Biologiques, Montréal, Quebec, Canada; Société Duvetnor Ltée, Rivière-du-Loup, Quebec, Canada; Université du Québec à Rimouski, Département de Biologie, Chimie et Géographie, Rimouski, Quebec, Canada; Memorial University of Newfoundland, Department of Biology, St. John’s, Newfoundland and Labrador, Canada; University of Prince Edward Island, Department of Pathology and Microbiology, Charlottetown, Prince Edward Island, Canada; Canadian Wildlife Health Cooperative, Charlottetown, Prince Edward Island, Canada; Université de Montréal, Faculté de médecine vétérinaire, Centre québécois sur la santé des animaux sauvages, Montréal, Quebec, Canada; Environment and Climate Change Canada, Wildlife and Landscape Science Directorate, Mount Pearl, Newfoundland and Labrador, Canada; Yale University, Department of Ecology and Evolutionary Biology, New Haven, Connecticut, Canada; Environment and Climate Change Canada, Canadian Wildlife Service, Mount Pearl, Newfoundland and Labrador, Canada

**Keywords:** HPAI, bird flu, wild bird, waterfowl, mass mortality, mortality assessment, disease

## Abstract

H5N1 clade 2.3.4.4b viruses have caused significant mortality events in various wild bird species across Europe, North America, South America, and Africa. In North America, the largest impacts on wild birds have been in eastern Canada, where over 40,391 wild birds were reported to have died from highly pathogenic avian influenza (HPAI) between April and September 2022. In the year following, we applied previously established methods to quantify total reported mortality in eastern Canada for a full year October 2022, to September 2023. In this study, we (i) document the spatial, temporal and taxonomic patterns of wild bird mortality in the 12 months that followed the mass mortality event in the summer of 2022 and (ii) quantify the observed differences in mortality across the breeding season (April to September) of 2022 and 2023. In eastern Canada, there was high uncertainty about whether 2023 would bring another year of devastating HPAI-linked mortalities. Mortalities in the breeding season were 93% lower in 2023 compared to 2022 but encompassed a more taxonomically diverse array of species. We found that mortalities in the fall and winter (non-breeding season) were dominated by waterfowl, while mortalities during the spring and summer (breeding season) were dominated by seabirds. Due to a low prevalence of HPAI among the subset of tested birds, we refrained from broadly attributing reported mortalities in 2023 to HPAI. However, our analysis did identify three notable mortality events linked to HPAI, involving at least 1,646 Greater Snow Geese, 232 Canada Geese, and 212 Northern Gannets.

This study emphasizes the ongoing need for H5NX surveillance and mortality assessments as the patterns of mortality in wild populations continue to change.

## Introduction

The impact of highly pathogenic avian influenza (HPAI) clade 2.3.4.4b H5N1 viruses and their derived reassortants (H5Nx) on both wild and domestic birds in recent years has been unprecedented (Careen et al., 2024; Lane et al., 2023; Ramey et al., 2022). This virus has also been responsible for disease and mortality events in an unprecedented number of wild bird species worldwide (Avery-Gomm et al., 2024; Leguia et al., 2023; Molini et al., 2023; Rijks et al., 2022). Another concerning development with this virus has been the positive detection in a wide variety of marine mammals and terrestrial mesopredators (Alkie et al., 2023; Plaza et al., 2024).

In North America, the first confirmed case of HPAI clade 2.3.4.4b H5N1 in a wild bird was detected in a Great Black-backed Gull (*Larus marinus*) in late 2021 in Newfoundland, Canada (Caliendo et al., 2022). Since then, mass mortality events in seabirds (e.g., Pelecaniformes, Suliformes, and Charadriiformes), sea ducks (e.g., American Common Eider; *Somateria mollissima*), and geese (Anseriformes; waterfowl) have been documented (Avery-Gomm et al., 2024; Giacinti et al., 2023). To date, the largest mortalities associated with HPAI in North America have been in eastern Canada, where at least 40,391 wild birds died during the 2022 HPAI outbreak between April and September 2022 (Avery-Gomm et al., 2024). The scale of mass mortality that occurred among seabirds and sea ducks was unanticipated and coincided with similar mass mortalities in Europe (European Food Safety Authority et al., 2022; Pearce-Higgins et al., 2022).

The rapidly evolving nature of HPAI viruses has complicated response and monitoring efforts, as its impacts are challenging to predict, prepare for, and manage (Ramey et al., 2022). In eastern Canada, there was high uncertainty about whether 2023 would bring another year of devastating HPAI-linked mortalities. In addition to increasing disease surveillance through Canada’s Interagency Surveillance Program for Avian Influenza Viruses in Wild Birds, Environment and Climate Change Canada (ECCC), the federal wildlife agency responsible for the conservation and management of migratory birds, opted to prepare for another significant mortality event in eastern Canada.

Using methods developed from Avery-Gomm et al. (2024), to document mortality in the spring/summer of 2022, we collated mortality reports across eastern Canada for 12 months spanning October 2022 to September 2023. Unlike Avery-Gomm et al. (2024), who sought to assess HPAI-linked mortality only, this study reports on complete mortality in wild birds regardless of the cause of death and uses avian influenza virus (AIV) screening data when available to draw inferences about the cause of death. The objectives of this study are to (i) document the spatial, temporal and taxonomic patterns of wild bird mortality in the 12 months that followed the spring/summer mass mortality events of 2022 described by Avery-Gomm et al. (2024), and (ii) quantify the observed differences in mortality across the breeding seasons (i.e., April and September) of 2022 and 2023.

## Materials and Methods

To the extent possible, the collation of mortality reports and data processing were aligned with the methods of Avery-Gomm et al. (2024). Accordingly, we defined eastern Canada as the provinces of Québec (QC), New Brunswick (NB), Nova Scotia (NS), Prince Edward Island (PE), and Newfoundland and Labrador (NL). We defined spring/summer as April 1 – September 30, which spans the breeding season for almost all birds in eastern Canada. To quantify the differences in mortality across the breeding seasons across two years, we present data for i) the spring/summer of 2022 (the period reported in Avery-Gomm et al. 2024) and ii) the spring/summer of 2023. In addition, we report mortalities for the intervening period (October 2022 – March 2023), hereafter referred to as the fall/winter period. Deviations from the methods described by Avery-Gomm et al. (2024) included an increased effort to survey beaches in the spring/summer of 2023, the inclusion of the complete mortality dataset (i.e., not only HPAI-linked mortality), and no effort to characterize the age classes of any species using photos submitted to iNaturalist.

In the spring and summer of 2023, preparations involved allocating financial and human resources to survey breeding colonies (aerial and on foot) for impacted populations of seabirds, and to conduct beach surveys. The goal of the colony surveys was to assess the impact of the 2022 outbreak on breeding colonial seabirds and sea ducks and the goal of the beach surveys was to improve the assessment of mortality events if one should occur by providing information on the onset, duration, and scale of mortality, as well as improving access to fresh carcasses for testing, necropsy, and early confirmation of HPAI.

### Collation of wild bird mortality data

For the first period (spring/summer 2022) the complete mortality dataset, which includes all reported mortalities - not only those attributed to HPAI, was obtained from Avery-Gomm et al. (2024). For the subsequent 12 months (October 2022 to September 2023), we requested and collated the submission of mortality reports from the same data providers that contributed data in the summer of 2022. This included provincial, federal, Indigenous, and municipal government staff and databases, the Canadian Wildlife Health Cooperative (CWHC), NGOs, university researchers, and two citizen science platforms (eBird and iNaturalist). The details on how data from iNaturalist were obtained are available in the Supplementary material (Appendix S1). Observations of wild bird mortalities on seabird and sea duck colonies, that were visited by government biologists and university researchers, were obtained through direct solicitation. Among the colonies visited were all six Northern Gannet (*Morus bassanus*) colonies, previously impacted Common Eider colonies in the St. Lawrence Estuary and some Common Murre (*Uria aalge*) and Herring Gull (*Larus argentatus*) colonies which were previously impacted (e.g. Taylor et al. 2023). Dates and details regarding which colonies were visited in 2023 for Northern Gannet, Common Eider, and Common Murres can be found in the online data repository (Data S2, Data S3, and Data S4 respectively).

Mortalities reported by data providers were supplemented with an increased beach survey effort during the spring/summer of 2023. From March 10 to November 3, 2023, 42 beaches across NB (13), NS (13), PE (14), and the island of Newfoundland (NF; 2) were surveyed. Fifteen of the beaches were selected because they had a high number of reported mortalities in the spring/summer of 2022. These were surveyed at 7 to 14-day intervals whenever possible. Poor weather or logistical issues (i.e., travel, staff availability) resulted in a maximum of 1 month occurring between surveys for some beaches (Appendix S2; Table S1). Twenty-seven of the beaches were surveyed on an ad hoc basis in response to public reports of mortality or to increase coverage of the coastline surveyed and these were surveyed following the same protocol, but at less regular intervals (i.e., 1-5 visits throughout the spring/summer).

Beach surveys followed standardized protocols (as described in Wilhelm et al. 2009, Lucas et al. 2012); they were conducted by walking along the lowest wrack line (i.e., the line of stranded seaweed and debris closest to the water) while scanning side to side for the entire length of the beach and returning along the next wrack line above. For wide beaches with many wrack lines, surveyors walked in a zig-zag motion to increase the encounter rate of carcasses. Carcasses that were identified as suitable for necropsy were collected following the HPAI beach survey protocol (Appendix S2) and carcasses too decomposed for testing were flagged or disposed of to avoid duplicate counting during the next survey. The site name, date, and start and end point of each survey were recorded (Appendix S2; Table S1).

### Data processing

All reports of mortality, including mortalities observed on colonies and during beach surveys, included the date, observer name and contact information, source, location (coordinates and/or site name), species identified to the lowest possible taxonomic level, and number of mortalities per species. When coordinates were unavailable or not provided in reports, georeferencing based on site name was conducted manually using Google Earth Pro or automatically using the geocode function in the R package ggmap (Kahle and Wickham, 2013). For observations that could not be identified to the species level, less precise taxonomic assignments were used (e.g., unknown gull, unknown tern). All observations were assigned to one of the following species groups: seabirds (includes American Common Eider), waterfowl, waders (e.g., herons, egrets, cranes), shorebirds (e.g., sandpipers, plovers, avocets, oystercatchers, phalaropes), loons, landbirds, raptors, or unknown.

### Attributing causes of mortality

We chose not to make broad assumptions about the cause of death within our study region during our study period. Therefore, the complete mortality dataset presented in this paper includes all reports of mortality regardless of cause of death, HPAI test results, or whether species were presumed HPAI virus-positive by Avery-Gomm et al. (2024) in 2022, with birds of unknown species also included. As part of Canada’s Interagency Surveillance Program for Avian Influenza Viruses in Wild Birds, a subsample of sick and dead wild birds was tested for the HPAI virus. We only attribute HPAI as a cause of death under two instances: 1) sick and dead birds that tested positive for HPAI, and 2) notable mortality events, which we define as a cluster of ≥100 mortalities of a single species within a 4-week period in the same location (i.e., province), where the prevalence of HPAI among the subsample of tested birds was >50%. For notable mortality events that meet the 50% threshold, we attribute all of the mortalities in the notable mortality event to HPAI. We report the sum of the total reported mortalities across all notable mortality events that are attributable to HPAI, but this must be viewed as a very conservative estimate of HPAI-linked mortality.

### Double-count analysis

Avery-Gomm et al. (2024) present a method for identifying instances within a dataset where records of a particular species reported in close proximity and time might suggest that the observation was reported more than once (i.e., double counts). This method was used to conduct a double-count analysis on the complete mortality dataset (April 2022 to September 2023). Following Avery-Gomm et al. (2024), we present results which exclude the smaller of two records that are identified as potential double counts because they belonged to the same species and were observed within 1 km and 1 day of each other, unless they were reported by the same person to the same source (Scenario B). Mortalities with less specific taxonomic assignments were handled as described in Avery-Gomm et al., (2024). For example, similar-looking gull species groups, that could be misidentified for one another were grouped with all reports of unknown gulls and included in the double-count analysis as ’Gulls’ (i.e., Gulls represents white-headed gulls; Herring, Great Black-backed, Lesser Black-backed (*Larus fuscus*), Ring-billed (*Larus delawarensis*), Iceland (*Larus glaucoides*), Glaucous (*Larus hyperboreus*), and unknown gulls). In the analysis, we also similarly defined Cormorants and Terns following Avery-Gomm et al. (2024). Other reports of less specific taxonomic assignments, that could not be grouped or considered interchangeable, were excluded from the analysis. The excluded reports were added to the double-count corrected dataset post-analysis.

### Species-specific mortality estimates and temporal comparisons

For the 12 months following the mass mortality events of spring/summer 2022 (i.e., October 2022 to September 2023) we present spatial, temporal, and taxonomic patterns of reported wild bird mortality using the complete mortality dataset with records identified as double counts excluded. The summed mortality numbers presented are minimum estimates, as we did not correct for birds that died that went unrecorded (e.g., areas not surveyed, birds not detected, birds not reported, birds lost at sea). Observed differences in species-specific mortality between the breeding seasons (e.g., April and September) of 2022 and 2023 are quantified using the complete, double-count corrected mortality dataset.

## Results

### Wild bird mortalities from October 1, 2022, to September 30, 2023

Through this mortality assessment, mortality reports were collated from four primary sources. Direct reports made by ECCC staff contributed 30.5% of reported mortalities (28% of these reports were documented during beach surveys). Provincial government reports contributed 24% of reported mortalities. As part of Canada’s Interagency Surveillance Program for AIV in Wild Birds, the Canadian Wildlife Health Cooperative contributed reports of sick and dead birds that accounted for 23% of the total reported mortalities. Data exported from iNaturalist and eBird contributed 17% of the reported mortalities. University researchers who work closely with wild bird populations and visit breeding colonies during the spring and summer contributed 4.5% of the reported mortalities. Parks Canada contributed 0.2% of reported mortalities. NGOs (e.g., NatureNB, Birds Canada, The Rock Wildlife Rescue) and Indigenous government partners were contacted in the region but very few had observations of mortality to report and cumulatively contributed 0.8%.

In total, 7,314 wild bird mortalities were recorded in eastern Canada in the 12 months that followed the mass mortality events described by Avery-Gomm et al., (2024). After the double count analysis, we found that 6.8% of these mortalities were likely birds that were reported by more than one observer, to more than one source, or both. Consequently, these redundant reports were excluded, leading to 6,977 unique wild bird mortalities across the region. These reports represent 199 species and 17 less precise taxonomic assignments. Among the provinces, the highest number of mortalities were reported in QC (3,501) and NS (2,156), followed by NL (569), PE (373), and NB (378).

The largest proportion of mortality for the entire annual cycle (October 2022 to September 2023) occurred in waterfowl (2,439; 35.0%) and seabirds (2,208; 31.6%), with smaller numbers of landbirds (1,351; 19.4%), raptors (775; 11.1%), waders (80; 1.1%), loons (71; 1.0%), shorebirds (30; 0.4%), and unknown species (23; 0.4%). Over the entire period, six species made up 50% of mortalities: Greater Snow Goose (*Anser caerulescens atlanticus*, 1701; 25%), American Crow (*Corvus brachyrhynchos,* 508; 7%), Canada Goose (*Branta canadensis,* 497; 7%), Northern Gannet, 368; 5%), Great Shearwater (*Ardenna gravis*, 288; 4%), and Herring Gull (239; 3%). The majority of mortality reported in the fall/winter (October 2022 to March 2023) was waterfowl (59% of 3,594), whereas mortality in the spring/summer (April to September 2023) consisted mostly of seabirds (47% of 3,383). A comprehensive breakdown of each group, species, and double-count corrected mortality can be found in Appendix S3.

### Notable mortality events

Across the entire study period, there were five notable mortality events, as defined as ≥100 mortalities in a province within a month: three in fall/winter and two in spring/summer. Out of the 6,977 mortalities in our complete double count corrected dataset, we directly attribute 302 mortalities to HPAI based on positive test results. In total, 2,076 mortalities were attributed to HPAI based on a high prevalence in the subsample of tested birds associated with those notable mortality events (Figure 1).

**Figure 1:**
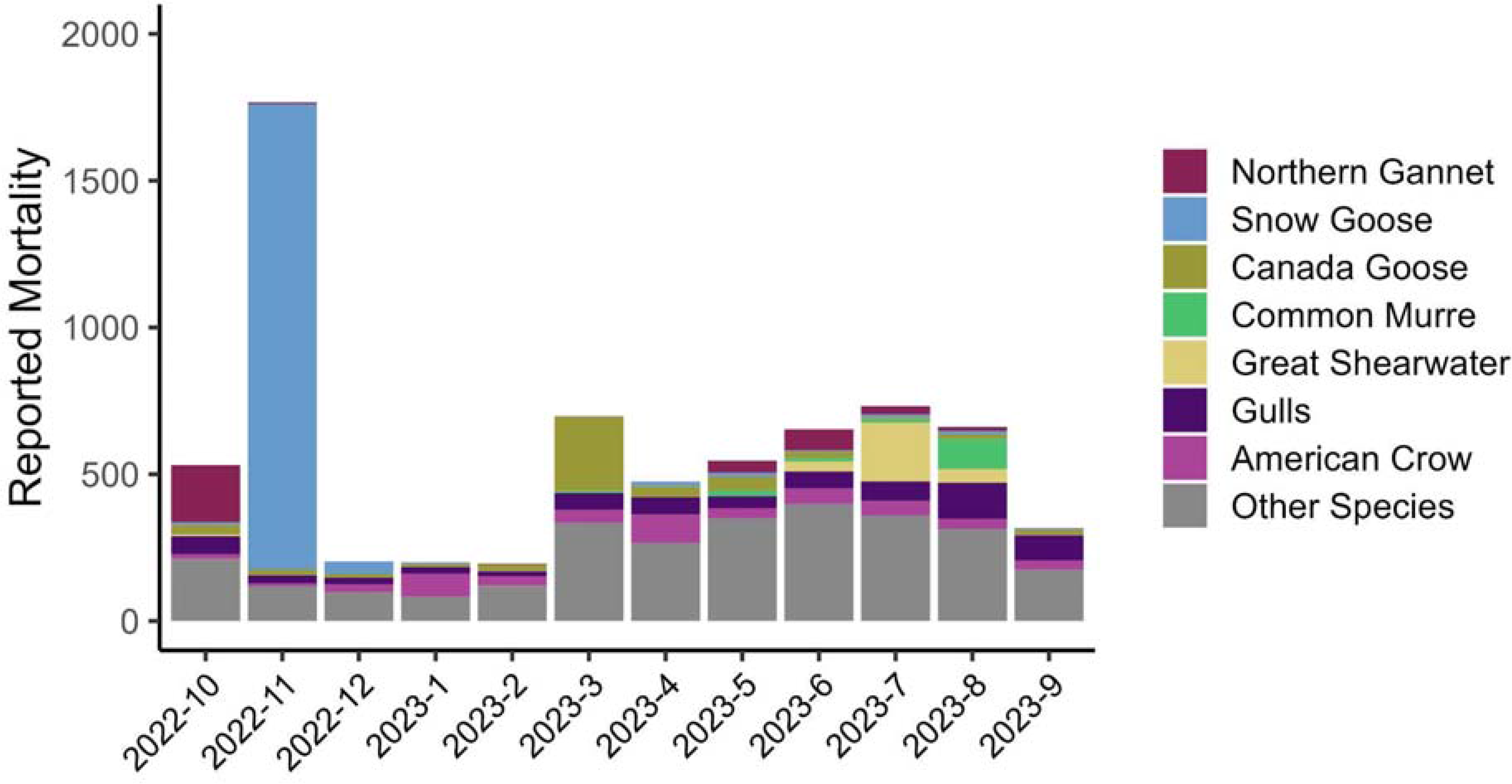
The taxonomic and temporal patterns of reported wild bird mortalities in the 12 months that followed the spring/summer mass mortality events of 2022 described by Avery-Gomm et al. (2024) in eastern Canada. Species with noteworthy levels of mortality or notable mortality events (i.e., ≥150 reports across the entire period) are coloured by species. ‘Other Species’ represents the remaining 209 species reported throughout the entire study period.

The three notable mortality events in the fall/winter included Northern Gannet, Greater Snow Goose, and Canada Goose (Figure 2A). In October 2022, a survey conducted in three study plots on Bonaventure Island, a breeding colony in QC, reported at least 212 dead Northern Gannets. These mortalities were associated with HPAI-linked mortality during the summer of 2022 at this location, which started in May 2022 and continued beyond the end of the study period described by Avery-Gomm et al. (2024). The second notable mortality event occurred in November 2022, involving 1,643 Greater Snow Geese in southern QC. A subset of Snow Geese carcasses was collected (68), and HPAI was detected in 92%. The third notable mortality event occurred on PE between late February and mid-March and involved 221 Canada Geese. HPAI was also confirmed in a subset of carcasses that were tested from this event (73% of 30 collected). Over half (58%) of the mortalities during the fall/winter were associated with these three mortality events (2,076 out of 3,594).

**Figure 2:**
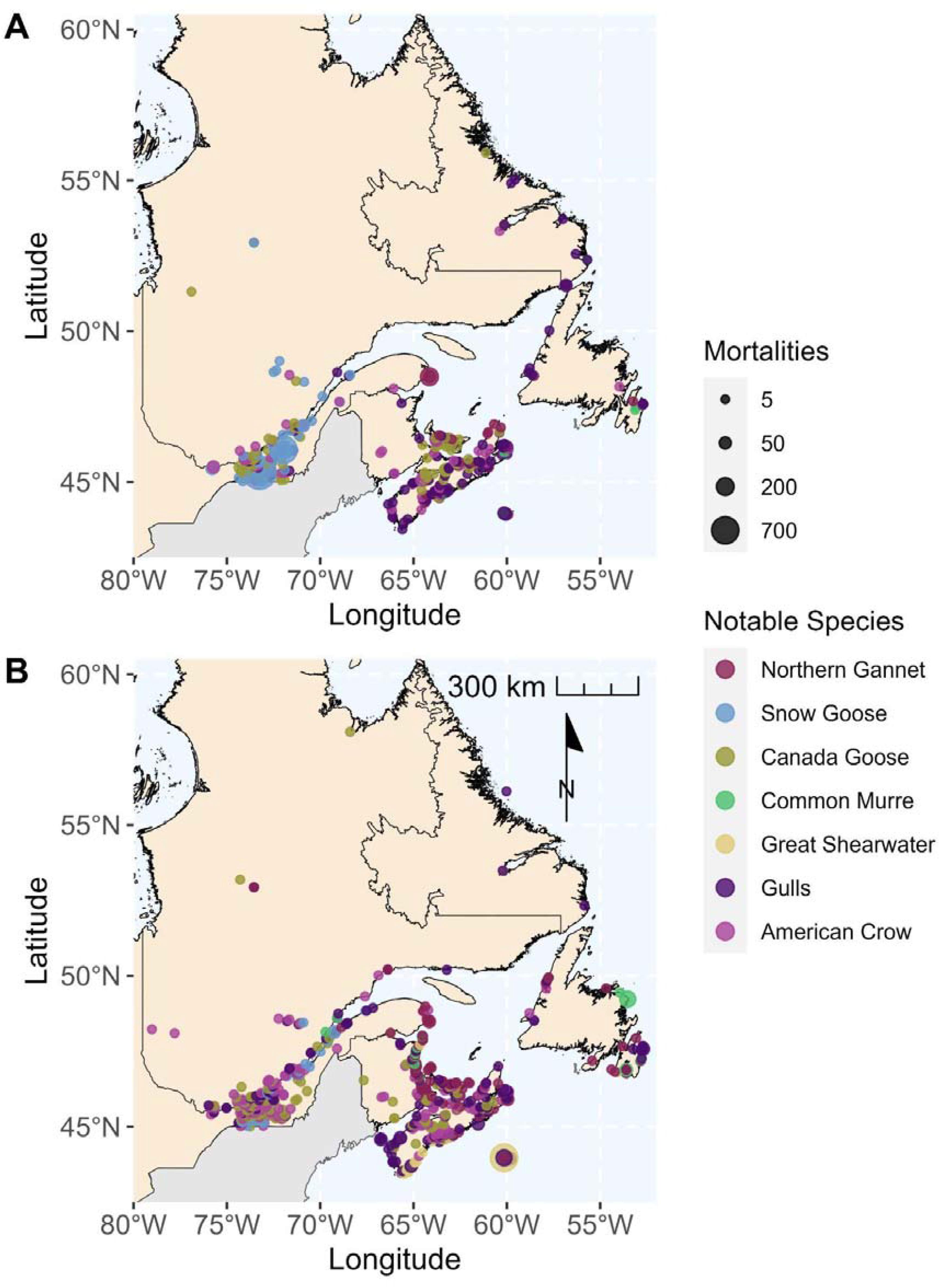
Spatial, temporal, and taxonomic pattern of reported mortalities in the 12 months that followed the spring/summer mass mortality events of 2022 described by Avery-Gomm et al., (2024) in eastern Canada during the (A) fall/winter period (October 2022 to March 2023) and (B) spring/summer period (April 2023 to September 2023). Only species with notable mortality events or noteworthy levels of mortality discussed in the text are shown.

The two notable spring/summer mortality events involved Great Shearwater and Common Murre (Figure 2B). In June and July 2023, highly emaciated Great Shearwaters washed ashore along the southern coast of NS. For example, a beach survey conducted on Sable Island, NS in July reported 132 dead Great Shearwater. Of those collected and sent for testing (7), none tested positive for the HPAI virus. In August 2023, at least 100 Common Murre washed ashore in Cape Freels, NL. Of the carcasses sent for testing (10), none tested positive for the HPAI virus.

Interestingly, following a positive detection of HPAI in a dead Common Tern (*Sterna hirundo*) on North Brother Island, NS in June, elevated levels of mortality were reported among tern chicks (> 25), including Roseate Tern (*Sterna dougallii*), an endangered species under the Species at Risk Act in Canada, that were also observed exhibiting neurological symptoms typical of HPAI. A subset of tern chicks (11) was collected and sent for testing and necropsy but none tested positive for HPAI. None of these mortalities were attributed to HPAI, other than the initial dead adult Common Tern tested.

Noteworthy mortality levels were also observed in American Crows (508) and Gulls (557) but were mostly reported as individual cases or in small numbers throughout the study period rather than as a single event (Figure 2B). HPAI was detected in both groups, but prevalence was relatively low (e.g., below the 50% threshold required to assume all mortalities were attributable to HPAI) and varied between seasons. During the fall/winter, 210 American Crows were reported (35% of 115 tested were positive) and during the spring/summer, an additional 298 American Crows were reported (11% of 98 tested were positive). During the fall/winter, 206 Gulls were reported (22% of 54 tested were positive). During the spring/summer of 2023, 434 Gulls were reported (19% of 66 tested were positive), including the only two HPAI virus detections from seabird breeding colonies (1 adult Herring Gull on Country Island, NS and 1 adult Herring Gull on Gull Island, NL).

### Interannual comparison of breeding season mortality

In the spring/summer of 2022, 44,595 unique wild bird mortalities were reported in the complete double-count corrected dataset. In the spring/summer of 2023, only 3,383 mortalities were reported, representing a 93% reduction in mortalities from the previous year. We attribute only 61 mortalities to HPAI (i.e., only those birds that were positive) in the spring/summer of 2023, compared to Avery-Gomm et al. (2024) who attributed 40,391 mortalities to HPAI during the same period in 2022. However, it is important to note the different approaches in how mortality was attributed, so these results should not be interpreted as a 99% reduction in HPAI among wild birds across the two years.

In 2022, 44% of mortalities during that period were associated with seabird breeding colonies, which was driven in large part by mass mortalities at four large Northern Gannet colonies in QC and NL. Correspondingly, the majority of mortalities were reported in these two provinces (19,158 and 16,661, respectively). In 2023, only a small percentage of mortalities were reported in colonies (148; 4.3% of spring/summer reported mortalities). The largest proportion of mortalities came from NS (1313; 39% of mortalities reported in 2023) closely followed by QC (1228; 36%), and relatively few mortalities were reported in NL (409, 12%).

Compared to 2022, the reported mortalities during the spring/summer of 2023 encompassed a more taxonomically diverse array of species. For instance, during this period in 2022, the majority of mortalities were seabirds and sea ducks (92%) and mortality among other species groups was limited. In stark contrast, although mortalities during the same period in 2023 were still largely seabirds (47%), more mortalities were reported in landbirds, waders, and raptors (Table 1).

**Table 1:** Interannual comparison of reported mortalities for each species group in the spring/summer period (April 1 to September 30). Our complete mortality dataset, with double counts removed, includes all reports of mortality regardless of the cause of death, HPAI test results, or whether species were presumed positive by Avery-Gomm et al., (2024) in 2022. Number of species included in each group is denoted in parentheses beside the totals. American Common Eider are included in seabirds.

| Group | 2022 |  | 2023 |  | Total Ratio of<br>2023 over 2022 |
| --- | --- | --- | --- | --- | --- |
|  | Total Birds<br>(# of<br>Species) | Percent of<br>Total | Total Birds<br>(# of<br>Species) | Percent of<br>Total |  |
| Seabirds | 40,911 (32) | 91.74% | 1,599 (35) | 47.27% | 0.04 |
| Landbirds | 441 (69) | 0.99% | 880 (92) | 26.09% | 2.0 |
| Waterfowl | 299 (21) | 0.67% | 324 (18) | 9.61% | 1.1 |
| Raptors | 108 (17) | 0.24% | 408 (24) | 12.10% | 3.8 |
| Loons | 39 (3) | 0.09% | 56 (3) | 1.66% | 1.4 |
| Waders | 23 (7) | 0.05% | 71 (6) | 2.10% | 3.1 |
| Shorebirds | 12 (8) | 0.03% | 24 (9) | 0.71% | 2.0 |
| Unknown<br>Bird | 2,762 | 6.19% | 21 | 0.62% | 0.008 |
| **Total** | **44,595** | **-** | **3,383** | **-** | **0.07** |

In 2022, most mortalities were dominated by only two species: Northern Gannet (26,199) and Common Murre (8,167), which constitute 77% of the overall reported mortality. In 2023, no single species represented more than 10% of overall mortality (Appendix S3). The two species with the highest number of mortalities in 2023 include the American Crow (9%) and Great Shearwater (8%) followed by Northern Gannet, Herring Gull, unknown gulls, Canada Goose, and Common Murre each representing between 4-5% of the overall mortality. The remaining 192 species each represent less than 4% of the total mortality.

Mortality among Northern Gannets and Common Murres in the breeding season of 2023 (156 and 145 respectively) was substantially lower than in 2022 (26,199 and 8,167 respectively). Although there was no mass mortality of either species in 2023, HPAI was detected in 3 gannets in June (2 in NB, 1 in QC) and not detected in any Common Murres collected for testing. In contrast, reported mortality for American Crows and Great Shearwaters was significantly higher in 2023 (298 and 282 respectively) than it was in 2022 (99 and 34 respectively).

## Discussion

In this study, we documented a substantial decline in reports of sick and dead birds in the 12 months that followed the largest HPAI-linked mortality event in North America (eastern Canada, spring/summer 2022; Avery-Gomm et al. 2024). A third of these mortalities can be attributed to HPAI, mostly associated with notable mortality events in late autumn and early spring involving Greater Snow Goose and Canada Goose - waterfowl species that did not suffer large mortalities during the 2022 breeding season in eastern Canada (Avery-Gomm et al., 2024). In the summer of 2023, we noted far fewer mortalities reported in Northern Gannet, Common Murre and Common Eider, the three species most affected in the summer of 2022, a pattern that is consistent with findings from the UK (Tremlett et al., 2024). Reported mortality was higher among landbirds, raptors and waterfowl during 2023 than in 2022. Spatially, mortalities concentrated around the St. Lawrence Estuary, and the coasts of NS and PE, with relatively few reports from elsewhere.

Assessing mortality based on collated reports from numerous sources, across a large study area comes with a set of caveats and precautions to consider when interpreting the data. Many of these are described in detail by Avery-Gomm et al. (2024), and equally apply to this study. Namely, mortalities should be viewed as a conservative estimate of true mortality because it is likely that many bird carcasses were not observed or reported, even in areas where survey effort was intensive. We feel confident that no mass mortality events were overlooked by our dataset, however, detection and reporting of mortality is likely to be lower in areas with lower human population density and at sea. The scope of this study differs from Avery-Gomm et al. (2024) in two key ways. First, we describe complete mortality rather than HPAI-linked mortality in 2023 because the prevalence of HPAI among the subset of tested dead birds was low. Although we attribute mortalities to HPAI if they were associated with a notable mortality event in which >50% of sampled dead birds tested positive, this likely underestimates HPAI-linked mortalities.

### Mortality in fall/winter

The largest notable HPAI-linked mortality events involved Greater Snow Goose (1,643) and Canada Goose (232). Generally, avian influenza viruses are common in waterfowl species, and cases are typically higher in the fall when large congregations of juveniles lacking immunity, mix with mature adults before migration (Arsnoe et al., 2011; Groepper et al., 2014). These pre-migration staging areas, combined with cooling temperatures in the fall (i.e., conditions favourable for the virus, Webster et al. 1992) likely facilitated the spread of the HPAI virus among Snow Geese in southern QC. The notable mortality event involving Canada Geese occurred later in the fall/winter period (March 2023), and this time of year birds can be more susceptible to infection due to colder winter temperatures, limited food availability, and generally lower body condition (Moon et al., 2007; Van Gils et al., 2007; Wight et al., 2024). The third notable mortality event involved at least 212 Northern Gannets reported on Bonaventure Island in October 2022. This mortality reflects the continuation of an ongoing HPAI-linked mortality event described by Avery-Gomm et al. (2024), which documented at least 3,611 mortalities between April 1 and September 30, 2022. Across eastern Canada, an estimated 26,193 gannet mortalities were reported over that period.

HPAI-linked mass mortality impacting wild bird populations is a very real concern that has been highlighted at the highest levels (CMS FAO Co-convened Scientific Task Force on Avian Influenza and Wild Birds, 2023). However, the significance of a mortality event depends on the ratio of mortalities to the size of the impacted population and the trend of the impacted population. Although Canada Goose populations are large globally (i.e., 7.6 million individuals; Partners in Flight 2021), populations are highly structured and have different management plans. The North Atlantic Population (NAP) breeds through Labrador, Québec, insular Newfoundland, and western Greenland, with wintering areas occurring primarily in coastal areas of southern Atlantic Canada and New England (Canadian Wildlife Service Waterfowl Committee, 2020; Mowbray et al., 2020). Historically, wintering populations existed farther south along the United States coast into the Carolinas but have since disappeared (Mowbray et al. 2020). Canada geese in the NAP have been formally managed as a separate population by the USFWS and CWS since 1996, with specific population and harvest management objectives (Atlantic Flyway Technical Section, 2008). The NAP of the Canada Goose has remained stable since 1990, estimated at approximately 52,500 breeding pairs (Canadian Wildlife Service Waterfowl Committee, 2020), but mixing with other Canada Goose populations during migration and harvest seasons makes it difficult to manage. We speculate that based on the timing and location of the notable mortality event (i.e., mid to late March in PE), most of the mortalities were likely NAP Canada geese. However, the magnitude of this specific mortality event (221 individuals) is not enough to cause management concerns.

In contrast to Canada Geese, all Snow Goose populations - which include the Lesser Snow Goose (*Chen caerulescens caerulescens*) and the Greater Snow Goose - have increased dramatically since the 1970s and 1930s, respectively, and are now considered overabundant (Lefebvre et al., 2017). The large Greater Snow Goose population (estimated at 900,000 in 2015; Lefebvre et al. 2017) means that our documented event involving 1,643 mortalities is not concerning from a conservation perspective. However, it is worth noting that notable mortalities in the hundreds or thousands have also been reported for Lesser Snow Geese in the Central Flyway and Pacific Flyway (Giacinti et al., 2023). Given the apparent susceptibility of all subspecies of Snow Geese to HPAI, efforts to collate mortality reports and disease surveillance data may be warranted if repeated mortality events from HPAI occur seasonally.

### Mortality in spring/summer 2023

There were only two notable mortality events during the spring and summer of 2023, both involving seabirds: Great Shearwater and Common Murre. None of the sampled birds of either species tested positive for the HPAI virus. Mortality events (sometimes called wrecks) for seabirds are not entirely uncommon. Past mortality events for seabirds, have been associated with oil spills or chronic oil pollution (Camphuysen, 1998; Wiese & Ryan, 2003; Wilhelm et al., 2007), bycatch in fishing gear (Davoren, 2007; Tasker et al., 2000) or starvation due to multiple factors including reduced food availability (Jones et al., 2018), summer heat waves (Piatt et al., 2020) or winter storms (Clairbaux et al., 2021; Diamond et al., 2020).

In 2022, Common Murre was the second most reported species and all mortalities were attributed to HPAI based on positive subsampled test results (Avery-Gomm et al. 2024). Therefore, it was unexpected when none of the birds tested from the notable mortality event in August 2023 were positive for HPAI. Carcasses collected from the mortality event indicated that the birds were in apparently good body condition and did not exhibit any signs of starvation or oiling and that bycatch in fishing gear was likely the cause of death (Ward, C. pers. comm.). In summer, inshore areas of eastern Newfoundland historically exhibit high rates of gillnet bycatch in murres and shearwaters (Hedd et al., 2016). From a conservation perspective, Common Murre populations in eastern Canada are large (estimated at 1.75 million breeding birds; Ainley et al. 2021). and the loss of 100 individuals of unknown age is not significant as a single event. However, this species is clearly under multiple pressures and there are concerns over the sustainability of these pressures, including a legal harvest (Cox et al., 2024).

None of the Great Shearwater mortalities in 2022 that were tested for HPAI were positive, and this was the result again in 2023. The occurrence of a notable mortality event for Great Shearwater is not entirely unexpected. Between 1993 and 2009, monthly beached bird surveys on Sable Island, NS by the Sable Island Institute consistently reported Great Shearwater carcasses mainly in June and July (1,304 carcasses total; Lucas et al. 2012) and low oiling rates reported to eliminate the possibility for oil pollution as a cause of death (Lucas et al., 2012). Over the same approximate period (1993-2013), at least twelve notable mortality events involving emaciated Great Shearwater were reported on the US East Coast (4,961 carcasses total; Haman et al. 2013, Robuck, A., pers comm).

Great Shearwaters are transequatorial migrants that travel from their breeding colonies in the southern hemisphere to their non-breeding grounds in the waters off the eastern coast of the United States and Canada where they forage during the summer months (Haman et al., 2013; Ronconi et al., 2010). The poor condition of these birds may reflect the cumulative stresses of evolving foraging conditions in their pre-migration staging areas off the coast of Argentina, a long migration, and inclement weather events once they arrive at their northern foraging grounds (Robuck, A., pers comm). In 2023, sea surface temperatures were among the highest on record (Cheng et al., 2024) and rapid oceanographic changes directly influence the abundance of forage fish and important prey species for marine top predators, like seabirds (Cairns, 1988; Harding et al., 2007). These factors together likely have contributed to the wrecking event of this long-distance migrant.

Despite not meeting our criteria for a notable mortality event, American Crow and Gulls (e.g., known species grouped with unidentified gulls) accounted for a high proportion of mortalities over the entire period. This mortality is difficult to distinguish from background rates because no historical mortality data in this manner exists. Crows and gulls are both scavengers, which is an established route for HPAI infection (Nemeth et al., 2023; van den Brand et al., 2018). While prevalence rates among crows collected for testing were high in the winter and mortality may therefore be attributed to HPAI, it was low in the summer. Prevalence rates were also low among gulls tested in both time periods.

Although gulls were among the breeding seabirds affected by HPAI in 2022 (Avery-Gomm et al. 2024), systematic daily monitoring on Kent Island, NB revealed order of magnitude lower mortality in 2023 (8 deaths, this study) compared to 2022 (>66 deaths; Taylor et al. 2023). Reduced mortality in the subsequent year may partially be a result of some degree of acquired immunity from previous exposure to HPAIs, or previous exposure to viruses with lower pathogenicity (e.g., H13 and H16; Huang et al. 2014, Benkaroun et al. 2016).

### Interannual comparison between 2022 and 2023 spring/summer mortalities

Mortalities in 2023 were represented by a more taxonomically diverse array of species than in 2022, which were dominated by seabirds and sea ducks. In 2023, we found that a larger proportion of mortalities were reported in landbirds, waders, and raptors, although seabirds still accounted for almost half of the reported mortalities. The most prominent contrast between breeding seasons is the remarkable decline in reported mortalities, dropping from 40,911 in 2022 to 1,599 in 2023. Several factors could explain this decline, including acquired immunity among seabird populations, genetic changes in circulating HPAI viruses, little to no presence of HPAI on the landscape, or any combination of these factors.

Before the arrival of HPAI H5N1 clade 2.3.4.4b in late 2021, wild birds in Canada were naïve to this particular clade of HPAIVs. Although some species (e.g., gulls, murres) may have had some immunity from past infection with low pathogenic avian influenza viruses (LPAIVs) that could have provided some protection against severe disease (Huang et al., 2013; Wille et al., 2014), naivety to HPAI was the likely cause of mass mortality experienced in 2022. In 2023, many wild bird populations across North America were no longer naïve to HPAI viruses. Those that survived infection in 2022, particularly Northern Gannets, Common Murres and Common Eiders, likely retained some level of immune protection in the year that followed (J. Provencher and J. Giacinti, unpublished data) which we posit explains the lower reported mortalities. Additionally, after more than a year of circulation across the continent, the ancestral virus responsible for high rates of mortality in 2022 (e.g., wholly Eurasian strain), has given rise to many reassortant HPAI viruses (Alkie et al. 2023, Giacinti et al. 2023, Signore et al. in prep). The extensive circulation of these genetically distinct reassortants, along with pre-existing immunity to HPAI, resulted in vastly different patterns of transmission, disease, and mortality than what occurred in 2022. Questions about how long immunity acquired from infection with the wholly Eurasian HPAI virus in 2022 will last and how this will protect seabirds and sea ducks in eastern Canada against severe outcomes from reassortant HPAI viruses remain.

### Role of Beached Bird Surveys in reporting mortalities

Due to HPAI, mortalities may be elevated compared to background rates, even when no notable mortality events occur. However, this is challenging to assess without historical baseline mortality data across the region. Systematic Beached Bird Surveys (BBS) provide the most robust way to assess long-term trends in seabird and sea duck mortalities (Lucas et al. 2012) but require significant time and effort. Beached bird survey programs previously existed across eastern Canada to diagnose, quantify, and audit the impacts of oil on seabirds, but have been scaled back since oiling rates declined in the 2000s (Wilhelm et al. 2009). Future studies could assess whether reported mortalities in the spring/summer of 2023 are above background rates, using historical data from existing surveys in NL and NS.

Another advantage of BBS is that they can increase access to fresh carcasses for HPAI testing and necropsies, and help detect the onset, duration and magnitude of mortality events, should they occur. BBS were implemented in eastern Canada in 2023, as recommended in Avery-Gomm et al., (2024), however no HPAI-linked mass mortality events occurred during this time. Although BBS conducted in this study helped document almost a quarter of the mortalities reported, very few of the birds reported on beaches were fresh enough for collection. Given the unpredictable nature of mass mortality events, we suggest a balanced program that targets a smaller number of beaches with high catchment rates for long-term monitoring to establish a baseline for annual mortality, with the ability for response teams to implement additional surveys during potential outbreak years.

## Conclusion

Globally, tens of thousands of HPAI outbreak notifications involving wild birds have been reported since 2021 (Wille & Waldenström, 2023), and many of these have been associated with mass mortality events. In 2023, Eastern Canada did not experience mass mortality events on the same scale as occurred in 2022, but experiences from abroad suggest this may have been the exception rather than the rule. For example, mass mortality in seabirds has been reported in the United Kingdom and Scotland in both 2022 and 2023 (Black-headed Gulls *Chroicocephalus ridibundus*, Black-legged Kittiwakes *Rissa tridactyla* and Sandwich *Thalasseus sandvicensis*, Common, and Arctic Terns *Sterna paradisaea*; Tremlett et al. 2024). In North America, the populations of many bird species are in decline (Rosenberg et al., 2019) and are under multiple pressures from the cumulative effects of human activities and climate change (Bateman et al., 2020; Phillips et al., 2023). Despite low mortalities in Canada in 2023, the H5Nx virus is a significant concern at a continental and global scale, and “*understanding HPAI impacts requires research and better data from outbreak situations, improved and standardized recording systems for wildlife settings, greater virus phylogenetic analyses, good population monitoring and research efforts”* (CMS FAO Co-convened Scientific Task Force on Avian Influenza and Wild Birds, 2023).

## Acknowledgements

Most reported mortalities can be traced back to engaged members of the public, and we offer our sincere thanks to all individuals who shared their observations. Numerous individuals and organizations have been instrumental in completing this research, and their collective contributions have played a pivotal role in shaping the research outcomes (Appendix S4).

Special thanks to Anna Robuck, Andrew Kennedy, Carina Gjerdrum, Catherine Soos, Georgia Taylor, Gretchen McPhail, Heather Major, Mike Brown, Jennifer F. Provencher, Jennifer Rock, Julie McKnight, Kate Bond, Mark Maddox, Meghan Baker, Raphaël A. Lavoie, Regina Wells, Shawn R. Craik, Sydney Collins, Yohannes Berhanes, and Zoe Lucas. Funding for this research was provided by Environment and Climate Change Canada’s Science and Technology Division.

## Author contributions

Tabatha L. Cormier: Conceptualization, Data curation, Formal analysis, Investigation, Methodology, Software, Validation, Visualization, Writing - original draft, Writing - review & editing. Tatsiana Barychka: Conceptualization, Data curation, Formal analysis, Software, Validation, Visualization, Writing - review & editing. Matthieu Beaumont: Data curation, Investigation, Methodology. Tori V. Burt: Investigation, Methodology, Writing - review & editing. Matthew D English: Data curation, Investigation, Methodology, Supervision, Writing - review & editing. Jolene A. Giacinti: Data curation, Investigation, Methodology, Writing - review & editing. Jean-François Giroux: Data curation, Investigation, Methodology, Writing - review & editing. Magella Guillemette: Investigation, Methodology, Resources, Supervision. Kathryn E. Hargan: Resources, Supervision, Writing - review & editing. Megan Jones: Investigation, Methodology, Resources, Supervision. Stéphane Lair: Investigation, Methodology, Resources, Supervision, Writing - review & editing. Andrew S. Lang: Investigation, Methodology, Resources, Supervision, Writing - review & editing. Christine Lepage: Data curation, Investigation, Methodology, Writing - review & editing. William A. Montevecchi: Investigation, Methodology, Resources, Supervision, Writing - review & editing. Ishraq Rahman: Investigation, Methodology, Writing - review & editing. Jean-François Rail: Investigation, Methodology, Writing - review & editing. Gregory J. Robertson: Writing - review & editing. Robert A. Ronconi: Conceptualization, Investigation, Methodology, Resources, Writing - review & editing. Yannick Seyer: Data curation, Investigation, Methodology, Writing - review & editing. Liam U. Taylor: Investigation, Methodology, Writing - review & editing. Christopher R. E. Ward: Investigation, Methodology, Writing - review & editing. Jordan Wight: Investigation, Methodology, Writing - original draft, Writing - review & editing. Sabina I. Wilhelm: Investigation, Methodology, Writing - review & editing. Stephanie Avery-Gomm: Conceptualization, Funding acquisition, Project administration, Resources, Supervision, Validation, Writing - original draft, Writing - review & editing.

## Data availability

The complete mortality dataset (Data S1), the code to reproduce the double count analysis, and colony survey information (Data S2, S3, S4) will be published upon acceptance of the manuscript. The following DOI has been reserved for this purpose: https://doi.org/10.6084/m9.figshare.25492693.

## Notes

https://doi.org/10.6084/m9.figshare.25492693.

### Competing Interest Statement

The authors have declared no competing interest.

## References

Ainley, D. G., Nettleship, D. N., & Storey, A. E. (2021). Common Murre (Uria aalge), version 2.0. In Birds of the World (S. M. Billerman, P. G. Rodewald, and B. K. Keeney, editors). Cornell Lab of Ornithology. 10.2173/bow.commur.02

Alkie, T. N., Byrne, A. M. P., Jones, M. E. B., Mollett, B. C., Bourque, L., Lung, O., James, J., Yason, C., Banyard, A. C., Sullivan, D., Signore, A. V., Lang, A. S., Baker, M., Dawe, B., Brown, I. H., & Berhane, Y. (2023). Recurring Trans-Atlantic Incursion of Clade 2.3.4.4b H5N1 Viruses by Long Distance Migratory Birds from Northern Europe to Canada in 2022/2023. Viruses, 15(9), 1836. 10.3390/v15091836

Arsnoe, D. M., Ip, H. S., & Owen, J. C. (2011). Influence of Body Condition on Influenza A Virus Infection in Mallard Ducks: Experimental Infection Data. PLoS ONE, 6(8), e22633. 10.1371/journal.pone.0022633

Atlantic Flyway Technical Section. (2008). Management Plan for the North Atlantic Population of Canada Geese. Canada Goose Committee.

Avery-Gomm, S., Barychka, T., English, M., Ronconi, R., Wilhelm, S. I., Rail, J.-F., Cormier, T., Beaumont, M., Bowser, C., Burt, T. V., Collins, S., Duffy, S., Giacinti, J. A., Gilliland, S., Giroux, J.-F., Gjerdrum, C., Guillemette, M., Hargan, K. E., Jones, M., … Wight, J. (2024). Wild bird mass mortalities in eastern Canada associated with the Highly Pathogenic Avian Influenza A(H5N1) virus, 2022 (p. 2024.01.05.574233). bioRxiv. 10.1101/2024.01.05.574233

Bateman, B. L., Wilsey, C., Taylor, L., Wu, J., LeBaron, G. S., & Langham, G. (2020). North American birds require mitigation and adaptation to reduce vulnerability to climate change. Conservation Science and Practice, 2(8), e242. 10.1111/csp2.242

Benkaroun, J., Shoham, D., Kroyer, A. N. K., Whitney, H., & Lang, A. S. (2016). Analysis of influenza A viruses from gulls: An evaluation of inter-regional movements and interactions with other avian and mammalian influenza A viruses. Cogent Biology, 2(1), 1234957. 10.1080/23312025.2016.1234957

Cairns, D. K. (1988). Seabirds as Indicators of Marine Food Supplies. Biological Oceanography, 5(4), 261–271.

Caliendo, V., Lewis, N. S., Pohlmann, A., Baillie, S. R., Banyard, A. C., Beer, M., Brown, I. H., Fouchier, R. A. M., Hansen, R. D. E., Lameris, T. K., Lang, A. S., Laurendeau, S., Lung, O., Robertson, G., van der Jeugd, H., Alkie, T. N., Thorup, K., van Toor, M. L., Waldenström, J., Berhane, Y. (2022). Transatlantic spread of highly pathogenic avian influenza H5N1 by wild birds from Europe to North America in 2021. Scientific Reports, 12(1), 11729. 10.1038/s41598-022-13447-z

Camphuysen, C. J. (1998). Beached bird surveys indicate decline in chronic oil pollution in the North Sea. Marine Pollution Bulletin, 36(7), 519–526. 10.1016/S0025-326X(98)80018-0

Canadian Wildlife Service Waterfowl Committee. (2020). CWS Migratory Birds Regulation Report. https://publications.gc.ca/collections/collection_2020/eccc/CW69-16-52-2019-eng.pdf

Careen, N. G., Collins, S. M., D’Entremont, K. J. N., Wight, J., Rahman, I., Hargan, K. E., Lang, A. S., & Montevecchi, W. (2024). Highly pathogenic avian influenza resulted in unprecedented reproductive failure and movement behaviour by Northern Gannets. Marine Ornithology, 52, 121–128.

Cheng, L., Abraham, J., Trenberth, K. E., Boyer, T., Mann, M. E., Zhu, J., Wang, F., Yu, F., Locarnini, R., Fasullo, J., Zheng, F., Li, Y., Zhang, B., Wan, L., Chen, X., Wang, D., Feng, L., Song, X., Liu, Y., … Lu, Y. (2024). New Record Ocean Temperatures and Related Climate Indicators in 2023. Advances in Atmospheric Sciences. 10.1007/s00376-024-3378-5

Clairbaux, M., Mathewson, P., Porter, W., Fort, J., Strøm, H., Moe, B., Fauchald, P., Descamps, S., Helgason, H. H., Bråthen, V. S., Merkel, B., Anker-Nilssen, T., Bringsvor, I. S., Chastel, O., Christensen-Dalsgaard, S., Danielsen, J., Daunt, F., Dehnhard, N., Erikstad, K. E., … Grémillet, D. (2021). North Atlantic winter cyclones starve seabirds. Current Biology, 31(17), 3964–3971.e3. 10.1016/j.cub.2021.06.059

CMS FAO Co-convened Scientific Task Force on Avian Influenza and Wild Birds. (2023). Scientific Task Force on Avian Influenza and Wild Birds statement on H5N1 high pathogenicity avian influenza in wild birds—Unprecedented conservation impacts and urgent needs (pp. 1–26). https://www.cms.int/sites/default/files/publication/avian_influenza_2023_aug.pdf

Cox, A. R., Roy, C., Hanson, A., & Robertson, G. J. (2024). Canadian murre harvest management in the face of uncertainty: A potential biological removal approach. The Journal of Wildlife Management, 88(4), e22573. 10.1002/jwmg.22573

Davoren, G. K. (2007). Effects of GillDNet Fishing on Marine Birds in a Biological Hotspot in the Northwest Atlantic. Conservation Biology, 21(4), 1032–1045. 10.1111/j.1523-1739.2007.00694.x

Diamond, A. W., Mcnair, D. B., Ellis, J. C., Rail, J.-F., Whidden, E. S., Kratter, A. W., Courchesne, S. J., Pokras, M. A., Wilhelm, S. I., Kress, S. W., Farnsworth, A., Iliff, M. J., Jennings, S. H., Brown, J. D., Ballard, J. R., Schweitzer, S. H., Okoniewski, J. C., Gallegos, J. B., & Stanton, J. D. (2020). Two Unprecedented Auk Wrecks in the Northwest Atlantic in Winter 2012/13. Marine Ornithology, 48, 185–204.

European Food Safety Authority, European Centre for Disease Prevention and Control, European Union Reference Laboratory for Avian Influenza, Adlhoch, C., Fusaro, A., Gonzales, J. L., Kuiken, T., Marangon, S., Niqueux, É., Staubach, C., Terregino, C., Guajardo, I. M., Chuzhakina, K., & Baldinelli, F. (2022). Avian influenza overview June – September 2022. EFSA Journal, 20(10). 10.2903/j.efsa.2022.7597

Giacinti, J. A., Signore, A. V., Jones, M. E. B., Bourque, L., Lair, S., Jardine, C., Stevens, B., Bollinger, T., Goldsmith, D., British Columbia Wildlife AIV Surveillance Program (BC WASPs), Pybus, M., Stasiak, I., Davis, R., Pople, N., Nituch, L., Brook, R. W., Ojkic, D., Massé, A., Dimitri-Masson, G., … Soos, C. (2023). Avian influenza viruses in wild birds in Canada following incursions of highly pathogenic H5N1 virus from Eurasia in 2021/2022. 10.1101/2023.11.23.565566

Groepper, S. R., DeLiberto, T. J., Vrtiska, M. P., Pedersen, K., Swafford, S. R., & Hygnstrom, S. E. (2014). Avian Influenza Virus Prevalence in Migratory Waterfowl in the United States, 2007–2009. Avian Diseases, 58(4), 531–540. 10.1637/10849-042214-Reg.1

Haman, K. H., Norton, T. M., Ronconi, R. A., Nemeth, N. M., Thomas, A. C., Courchesne, S. J., Segars, A., & Keel, M. K. (2013). Great Shearwater (Puffinus gravis) Mortality Events Along the Eastern Coast of the United States. Journal of Wildlife Diseases, 49(2), 235–245. 10.7589/2012-04-119

Harding, A., Piatt, J., & Schmutz, J. (2007). Seabird behavior as an indicator of food supplies: Sensitivity across the breeding season. Marine Ecology Progress Series, 352, 269–274. 10.3354/meps07072

Hedd, A., Regular, P. M., Wilhelm, S. I., Rail, J., Drolet, B., Fowler, M., Pekarik, C., & Robertson, G. J. (2016). Characterization of seabird bycatch in eastern Canadian waters, 1998–2011, assessed from onboard fisheries observer data. Aquatic Conservation: Marine and Freshwater Ecosystems, 26(3), 530–548. 10.1002/aqc.2551

Huang, Y., Wille, M., Benkaroun, J., Munro, H., Bond, A. L., Fifield, D. A., Robertson, G. J., Ojkic, D., Whitney, H., & Lang, A. S. (2014). Perpetuation and reassortment of gull influenza A viruses in Atlantic North America. Virology, 456–457, 353–363. 10.1016/j.virol.2014.04.009

Huang, Y., Wille, M., Dobbin, A., Robertson, G. J., Ryan, P., Ojkic, D., Whitney, H., & Lang, A. S. (2013). A 4-year study of avian influenza virus prevalence and subtype diversity in ducks of Newfoundland, Canada. Canadian Journal of Microbiology, 59(10), 701–708. 10.1139/cjm-2013-0507

Jones, T., Parrish, J. K., Peterson, W. T., Bjorkstedt, E. P., Bond, N. A., Ballance, L. T., Bowes, V., Hipfner, J. M., Burgess, H. K., Dolliver, J. E., Lindquist, K., Lindsey, J., Nevins, H. M., Robertson, R. R., Roletto, J., Wilson, L., Joyce, T., & Harvey, J. (2018). Massive Mortality of a Planktivorous Seabird in Response to a Marine Heatwave. Geophysical Research Letters, 45(7), 3193–3202. 10.1002/2017GL076164

Kahle, D. & Wickham, H. (2013). ggmap: Spatial Visualization with ggplot2. 5(1), 144–161.

Lane, J. V., Jeglinski, J. W. E., Avery-Gomm, S., Ballstaedt, E., Banyard, A. C., Barychka, T., Brown, I. H., Brugger, B., Burt, T. V., Careen, N., Castenschiold, J. H. F., Christensen-Dalsgaard, S., Clifford, S., Collins, S. M., Cunningham, E., Danielsen, J., Daunt, F., D’entremont, K. J. N., Doiron, P., … Votier, S. C. (2023). High pathogenicity avian influenza (H5N1) in Northern Gannets (Morus bassanus): Global spread, clinical signs and demographic consequences. *Ibis*, Early View. 10.1111/ibi.13275

Lefebvre, J., Gauthier, G., Giroux, J.-F., Reed, A., Reed, E. T., & Bélanger, L. (2017). The greater snow goose Anser caerulescens atlanticus: Managing an overabundant population. Ambio, 46(S2), 262–274. 10.1007/s13280-016-0887-1

Leguia, M., Garcia-Glaessner, A., Muñoz-Saavedra, B., Juarez, D., Barrera, P., Calvo-Mac, C., Jara, J., Silva, W., Ploog, K., Amaro, Lady, Colchao-Claux, P., Johnson, C. K., Uhart, M. M., Nelson, M. I., & Lescano, J. (2023). Highly pathogenic avian influenza A (H5N1) in marine mammals and seabirds in Peru. Nature Communications, 14(1), Article 1. 10.1038/s41467-023-41182-0

Lucas, Z., Horn, A., & Freedman, B. (2012). Beached Bird Surveys on Sable Island, Nova Scotia, 1993–2009, Show a Decline in the Incidence of Oiling. Proceedings of the Nova Scotian Institute of Science, 47, 91–129.

Molini, U., Yabe, J., Meki, I. K., Ouled Ahmed Ben Ali, H., Settypalli, T. B. K., Datta, S., Coetzee, L. M., Hamunyela, E., Khaiseb, S., Cattoli, G., Lamien, C. E., & Dundon, W. G. (2023). Highly pathogenic avian influenza H5N1 virus outbreak among Cape cormorants (Phalacrocorax capensis) in Namibia, 2022. Emerging Microbes & Infections, 12(1), 2167610. 10.1080/22221751.2023.2167610

Moon, J. A., Haukos, D. A., & Smith, L. M. (2007). Declining Body Condition of Northern Pintails Wintering in the Playa Lakes Region. The Journal of Wildlife Management, 71(1), 218–221. 10.2193/2005-596

Mowbray, T. B., Ely, C. R., Sedinger, J. S., & Trost, R. E. (2020). Canada Goose (Branta canadensis), version 1.0. In Birds of the World (P.G. Rodewald, Editor). Cornell Lab of Ornithology. 10.2173/bow.cangoo.01

Nemeth, N. M., Ruder, M. G., Poulson, R. L., Sargent, R., Breeding, S., Evans, M. N., Zimmerman, J., Hardman, R., Cunningham, M., Gibbs, S., & Stallknecht, D. E. (2023). Bald eagle mortality and nest failure due to clade 2.3.4.4 highly pathogenic H5N1 influenza a virus. Scientific Reports, 13(1), 191. 10.1038/s41598-023-27446-1

Partners in Flight. (2021). *Avian Conservation Assessment Database, version 2021.* [dataset]. http://pif.birdconservancy.org/ACAD

Pearce-Higgins, J. W., Humphreys, E. M., Burton, N. H. K., Atkinson, P. W., Pollock, C., Johnston, D. T., O’Hanlon, N. J., Balmer, D. E., Frost, T. M., Harris, S. J., & Baker, H. (2022). Highly pathogenic avian influenza in wild birds in the United Kingdom in 2022: Impacts, planning for future outbreaks, and conservation and research priorities.

Phillips, R. A., Fort, J., & Dias, M. P. (2023). Chapter 2—Conservation status and overview of threats to seabirds. In L. Young & E. VanderWerf (Eds.), Conservation of Marine Birds (pp. 33–56). Academic Press. 10.1016/B978-0-323-88539-3.00015-7

Piatt, J. F., Parrish, J. K., Renner, H. M., Schoen, S. K., Jones, T. T., Arimitsu, M. L., Kuletz, K. J., Bodenstein, B., García-Reyes, M., Duerr, R. S., Corcoran, R. M., Kaler, R. S. A., McChesney, G. J., Golightly, R. T., Coletti, H. A., Suryan, R. M., Burgess, H. K., Lindsey, J., Lindquist, K., … Sydeman, W. J. (2020). Extreme mortality and reproductive failure of common murres resulting from the northeast Pacific marine heatwave of 2014-2016. PLoS ONE, 15(1), e0226087. 10.1371/journal.pone.0226087

Plaza, P. I., Gamarra-Toledo, V., Euguí, J. R., & Lambertucci, S. A. (2024). Recent Changes in Patterns of Mammal Infection with Highly Pathogenic Avian Influenza A(H5N1) Virus Worldwide—Volume 30, Number 3—March 2024—Emerging Infectious Diseases journal—CDC. Emerging Infectious Diseases, 30(3), 444–452. 10.3201/eid3003.231098

Ramey, A. M., Hill, N. J., DeLiberto, T. J., Gibbs, S. E. J., Camille Hopkins, M., Lang, A. S., Poulson, R. L., Prosser, D. J., Sleeman, J. M., Stallknecht, D. E., & Wan, X.-F. (2022). Highly pathogenic avian influenza is an emerging disease threat to wild birds in North America. The Journal of Wildlife Management, 86(2), e22171. 10.1002/jwmg.22171

Rijks, J. M., Leopold, M. F., Kühn, S., In ’t Veld, R., Schenk, F., Brenninkmeijer, A., Lilipaly, S. J., Ballmann, M. Z., Kelder, L., de Jong, J. W., Courtens, W., Slaterus, R., Kleyheeg, E., Vreman, S., Kik, M. J. L., Gröne, A., Fouchier, R. A. M., Engelsma, M., de Jong, M. C. M., … Beerens, N. (2022). Mass Mortality Caused by Highly Pathogenic Influenza A(H5N1) Virus in Sandwich Terns, the Netherlands, 2022. Emerging Infectious Diseases, 28(12), 2538–2542. 10.3201/eid2812.221292

Ronconi, R., Koopman, H., McKinstry, C., Wong, S., & Westgate, A. (2010). Inter-annual variability in diet of non-breeding pelagic seabirds Puffinus spp. at migratory staging areas: Evidence from stable isotopes and fatty acids. Marine Ecology Progress Series, 419, 267–282. 10.3354/meps08860

Rosenberg, K. V., Dokter, A. M., Blancher, P. J., Sauer, J. R., Smith, A. C., Smith, P. A., Stanton, J. C., Panjabi, A., Helft, L., Parr, M., & Marra, P. P. (2019). Decline of the North American avifauna. Science, 366(6461), 120–124. 10.1126/science.aaw1313

Tasker, M., Camphuysen, C. J. (Kees), Cooper, J., Garthe, S., Montevecchi, W. A., & Blaber, S. J. M. (2000). The impacts of fishing on marine birds. ICES Journal of Marine Science, 57(3), 531–547. 10.1006/jmsc.2000.0714

Taylor, L. U., Ronconi, R. A., Spina, H. A., Jones, M. E. B., Ogbunugafor, C. B., & Ayala, A. J. (2023). Limited Outbreak of Highly Pathogenic Influenza A(H5N1) in Herring Gull Colony, Canada, 2022. Emerging Infectious Diseases, 29(10). 10.3201/eid2910.230536

Tremlett, C. J., Morley, N., & Wilson, L. J. (2024). UK seabird colony counts in 2023 following the 2021-22 outbreak of Highly Pathogenic Avian Influenza (76; RSPB Research Report). RSPB Centre for Conservation Science (RSPB). https://base-prod.rspb-prod.magnolia-platform.com/dam/jcr:7983cad1-03f7-4ab4-b22e-c56c9f02243b/RSPB%20HPAI%20seabird%20counts%20report_Feb24.pdf

van den Brand, J. M. A., Verhagen, J. H., Veldhuis Kroeze, E. J. B., van de Bildt, M. W. G., Bodewes, R., Herfst, S., Richard, M., Lexmond, P., Bestebroer, T. M., Fouchier, R. A. M., & Kuiken, T. (2018). Wild ducks excrete highly pathogenic avian influenza virus H5N8 (2014–2015) without clinical or pathological evidence of disease. Emerging Microbes & Infections, 7(1), 1–10. 10.1038/s41426-018-0070-9

Van Gils, J. A., Munster, V. J., Radersma, R., Liefhebber, D., Fouchier, R. A. M., & Klaassen, M. (2007). Hampered Foraging and Migratory Performance in Swans Infected with Low-Pathogenic Avian Influenza A Virus. PLoS ONE, 2(1), e184. 10.1371/journal.pone.0000184

Webster, R. G., Bean, W. J., Gorman, O. T., Chambers, T. M., & Kawaoka, Y. (1992). Evolution and ecology of influenza A viruses. Microbiological Reviews, 56(1), 152–179. 10.1128/mr.56.1.152-179.1992

Wiese, F. K., & Ryan, P. C. (2003). The extent of chronic marine oil pollution in southeastern Newfoundland waters assessed through beached bird surveys 1984–1999. Marine Pollution Bulletin, 46(9), 1090–1101. 10.1016/S0025-326X(03)00250-9

Wight, J., Rahman, I., Wallace, H. L., Cunningham, J. T., Roul, S., Robertson, G. J., Russell, R. S., Xu, W., Zhmendak, D., Alkie, T. N., Berhane, Y., Hargan, K. E., & Lang, A. S. (2024). Avian influenza virus circulation and immunity in a wild urban duck population prior to and during a highly pathogenic H5N1 outbreak. 10.1101/2024.02.22.581693

Wilhelm, S. I., Robertson, G. J., Ryan, P. C., Tobin, S. F., & Elliot, R. D. (2007). An integrated approach to monitor year-round chronic oil pollution in southeastern Newfoundland waters. Proceedings: Papers, 229.

Wilhelm, S. I., Robertson, G. J., Ryan, P. C., Tobin, S. F., & Elliot, R. D. (2009). Re-evaluating the use of beached bird oiling rates to assess long-term trends in chronic oil pollution. Marine Pollution Bulletin, 58(2), 249–255. 10.1016/j.marpolbul.2008.09.018

Wille, M., Huang, Y., Robertson, G. J., Ryan, P., Wilhelm, S. I., Fifield, D., Bond, A. L., Granter, A., Munro, H., Buxton, R., Jones, I. L., Fitzsimmons, M. G., Burke, C., Tranquilla, L. M., Rector, M., Takahashi, L., Kouwenberg, A.-L., Storey, A., Walsh, C., … Lang, A. S. (2014). Evaluation of Seabirds in Newfoundland and Labrador, Canada, as Hosts of Influenza A Viruses. Journal of Wildlife Diseases, 50(1), 98–103. 10.7589/2012-10-247

Wille, M., & Waldenström, J. (2023). Weathering the Storm of High Pathogenicity Avian Influenza in Waterbirds. Waterbirds, 46(1), 100–109. 10.1675/063.046.0113

