## Appendix S1 for "Seabird and sea duck mortalities were lower during the second breeding season in eastern Canada following the introduction of Highly Pathogenic Avian Influenza A H5Nx viruses"

**Appendix S1. Methods for downloading and extracting mortality reports from iNaturalist**

iNaturalist projects filter through all observations based on a set of required criteria outlined by the creator of the project. The methods used to download and extract iNaturalist observations of dead birds using iNaturalist projects was adopted from Avery-Gomm et al., (2024, Appendix S1), with a few exceptions which are detailed in this Appendix.

***Wild Bird Mortality in Eastern Canada iNaturalist Project***

We created an iNaturalist project “**Wild Bird Mortality in Eastern Canada 2023**” to identify any bird species that was annotated as “Dead” in the study area during the study window. The study area was Quebec, New Brunswick, Nova Scotia, Prince Edward Island, Newfoundland and Labrador, and included the French territory of Saint Pierre Miquelon. The project can be accessed at <https://www.inaturalist.org/projects/wild-bird-mortality-in-eastern-canada-2023>

As was highlighted by Bartalotta et al., (2023) and Taylor et al., (in prep), some observers do not annotate dead birds as dead, and as a result, their observations may be missed from the “Wild Bird Mortality in Eastern Canada 2023.” **To ensure that observations of prolific users were not missed,** we identified the five iNaturalist observers who contributed the most observations in the complete mortality dataset published by Avery-Gomm et al., (2024). These were *lewnanny_richardson, audree_b, ahebert, alexisgodin,* and *courtjcam.*

For these observers, we searched each user on the iNaturalist web browser and filtered their observations for **birds only** within our specified range for the 2023 tracker (**April 1, 2022 to September 30, 2023**). We examined the images for each observation to identify where birds were visibly dead. If an observation of a dead bird was not already annotated as "Dead”, we manually added the annotation in iNaturalist.

In 2022, Northern Gannets were a species with high mortality, to ensure all records of Northern Gannets were captured in the 2023 dataset, **all observations of gannets** (i.e., no annotation) within the study area and period were examined and manually marked as dead. After annotating the appropriate observations as dead, we proceeded to export the data from the project. 

**Exporting data** 
We used the export function to extract observations at the iNaturalist website: <https://www.inaturalist.org/observations/export>.

To export data from “**Wild Bird Mortality in Eastern Canada 2023**”, we used the following query field, and selected all the options under ‘Geo’ and ‘Taxon Extras.’ Unlike Avery-Gomm et al. 2024, we did not target specific avian taxa groups and did not filter based on exclusion terms describing non-HPAI related causes of death. Therefore, data from this project was exported for April 1, 2022, to September 30, 2023.

*Query quality_grade=any&identifications=any&projects[]=wild-bird-mortality-in-eastern-canada-2023 Columns*

*id, observed_on_string, observed_on, time_observed_at, time_zone, user_id, user_login, user_name, created_at, updated_at, quality_grade, license, url, image_url, sound_url, tag_list, description, num_identification_agreements, num_identification_disagreements, captive_cultivated, oauth_application_id, place_guess, latitude, longitude, positional_accuracy, private_place_guess, private_latitude, private_longitude, public_positional_accuracy, geoprivacy, taxon_geoprivacy, coordinates_obscured, positioning_method, positioning_device, place_town_name, place_county_name, place_state_name, place_country_name, place_admin1_name, place_admin2_name, species_guess, scientific_name, common_name, iconic_taxon_name, taxon_id, taxon_kingdom_name, taxon_phylum_name, taxon_subphylum_name, taxon_superclass_name, taxon_class_name, taxon_subclass_name, taxon_superorder_name, taxon_order_name, taxon_suborder_name, taxon_superfamily_name, taxon_family_name, taxon_subfamily_name, taxon_supertribe_name, taxon_tribe_name, taxon_subtribe_name, taxon_genus_name, taxon_genushybrid_name, taxon_species_name, taxon_hybrid_name, taxon_subspecies_name, taxon_variety_name, taxon_form_name*

**Post processing of data:**

The export included 1014 observations.

Where iNaturalist records had quality_grade = needs_id (n= 219), photos were downloaded and examined to verify species identification to the lowest possible level, and this information was added to the dataset. Finally, these records were merged with the master list of wild bird mortalities.

**References**

Avery-Gomm, S., T. Barychka, M. English, R. Ronconi, S. I. Wilhelm, J.-F. Rail, T. Cormier, M. Beaumont, C. Bowser, T. V. Burt, S. Collins, S. Duffy, J. A. Giacinti, S. Gilliland, J.-F. Giroux, C. Gjerdrum, M. Guillemette, K. E. Hargan, M. Jones, A. Kennedy, L. Kusalik, S. Lair, A. S. Lang, R. A. Lavoie, C. Lepage, G. McPhail, W. A. Montevecchi, G. J. Parsons, J. F. Provencher, I. Rahman, G. J. Robertson, Y. Seyer, C. Soos, C. R. E. Ward, R. Wells, and J. Wight. 2024, January 5. Wild bird mass mortalities in eastern Canada associated with the Highly Pathogenic Avian Influenza A(H5N1) virus, 2022. BioRxiv. <https://www.biorxiv.org/content/10.1101/2024.01.05.574233v1>
