## Appendix S2 for "Seabird and sea duck mortalities were lower during the second breeding season in eastern Canada following the introduction of Highly Pathogenic Avian Influenza A H5Nx viruses"

**Appendix S2. Beached Bird Survey protocol modified to monitor mortality associated with Highly Pathogenic Avian Influenza (HPAI)**

ECCC increased beach survey effort during the spring/summer of 2023 with the goal to better understand the onset, duration, and magnitude of potential mortality events. Beach surveys also improve access to fresh carcasses for necropsy and HPAI testing, and provide information on species composition and age structure during a given mortality event.

From March 10 to November 3, 2023, 42 beaches across NB (13), NS (13), PE (14), and the island of Newfoundland (NF; 2) were surveyed.

- **Fifteen** of the beaches were selected because they had a high number of dead birds in the spring/summer of 2022. These were surveyed at 7 to 14-day intervals whenever possible.
- **Twenty-seven** of the beaches were surveyed on an ad hoc basis in response to public reports of mortality or with the goal of increasing coverage of coastline surveyed and these were surveyed following the same protocol, but at less regular intervals (i.e., 1-5 visits throughout the spring/summer).

Beach surveys followed a modified version of the beach survey protocol “*Adopt-a-beach – Newfoundland and Labrador Beached Bird Survey Program: Surveyor’s guide”* which was itself adapted from Chardine and Pelly (1994).

### **1. Protocol for Beached Bird Surveys**

### **When**: Conduct your survey at 7–14 day intervals on the same stretch of beach. It is best to walk the beach at low tide or as the tide is receding. Sometimes inclement weather or personal circumstances do not allow for regular outings. Surveys need to be conducted by trained personnel.

**Where**: The best beaches are those that are easy to walk, not too steep, composed of sand/gravel or small cobble and that naturally accumulate driftwood and seaweeds. Walk one way along the wrack or flotsam line (i.e., the line of stranded seaweed and debris) closest to the water, scanning on each side, and return along the next wrack line above. Note that fresh carcasses may also be found close to the water line, whereas old carcasses may be found at the extreme high tide. If a beach is particularly deep, several sweeps may be required.

### **2. Filling out the Datasheets**

#### **a. Survey Information**

Always complete all boxes relevant to each beach visit (beach name, observer name, and the day, month and year the survey was conducted) even if no birds are found.

#### **b. Carcass information**

**Species**: It is important to try to identify each carcass to its lowest taxonomic level (i.e., species). The Beached Bird field guide will be especially useful to identify carcasses.

When it is not possible to identify a carcass to species because it is too heavily scavenged or decomposed, provide as much information as you can (e.g., unknown gull (UNGU) or unknown murre (UNMU)). If you have a digital camera, you may want to take a photo of the carcass to confirm the identification with an experienced person at a later date.

**Age and sex**: In some birds, it is relatively easy to differentiate between immature and adult birds (e.g., gulls) and between males and females (e.g., sea ducks). Referring to the Beached Bird field guide can help you record this information.

**Degree of scavenging:** Evaluate how much of the body is intact (i.e., not scavenged or missing). Indicate by **Yes or No** if more than half of the breast is present.

**Degree of oiling:** Record whether there is oil on the carcass by **Yes or No**

#### **c. Recording Live Wildlife in Area**

At the bottom of the datasheet, you will find a space to record live birds present in the area (i.e., on the beach or on the water inshore) during your survey. Please record the number of live birds and what species, as well as their behaviours, (e.g., foraging at sea, scavenging on dead birds, resting on water) and whether they are showing neurological symptoms (e.g., lethargic, head twitching).

Additional space is provided for any notes or other observations you wish to record while conducting your beached bird survey. Please see below for an example of a blank datasheet.

### **3. Dealing with Bird Carcasses**

When fresh carcasses are encountered, a sub-sample may be collected to test for HPAI. **Whether there is a need to collect carcasses for testing should be discussed with your local ECCC-CWS coordinator (see below) prior to conducting a survey**.

- If you find a fresh whole bird suitable for HPAI testing, collect the bird as follows:
  - Put on a pair of clean disposable gloves and mask
  - Place bird in clear plastic bag and place this bag in another clear plastic bag (i.e., double-bag each bird)
  - Write on plastic bag the date, location, that bird was found dead, and your name and contact information. Where possible, include CWHC submission form inside bag: <http://www.cwhc-rcsf.ca/forms/cwhc_atlantic_submission_form.pdf>
  - Place in cool area sheltered from predators
  - Contact the coordinator of this program (see below) to arrange for pickup

### **It is also important to clear the beach of remaining carcasses after every survey to ensure that you do not recount the same bird during your next survey; these birds can be placed in a single large bag (or more as required), double-bagged, and disposed of in accordance to Provincial agency guidelines. Alternatively, carcasses can be marked using spray paint or a plastic zip tie (on leg or wing) so that they are recognized during subsequent surveys and not counted as a new beached bird.** Note that in either case, if the number of carcasses exceeds the capacity for removal or marking, this should be reported immediately to your local ECCC-CWS coordinator.

**Table S1.** Dates of ECCC beach surveys in eastern Canada between October 1, 2022, and September 30, 2023. Sites that were chosen because they had a high number of dead birds reported in the spring/summer of 2022 are indicated with (*). Other beaches were surveyed on an ad hoc basis in response to public reports of mortality or with the goal of increasing coverage of coastline surveyed.

| **PROVINCE**  **Beach, Latitude and Longitude, Date Visited** | **Birds Found** |
| --- | --- |
| **NOVA SCOTIA** |  |
| Ballantyne's Cove (45.85879, -61.9174) | 2 |
| 2023-06-10 | 2 |
| Big Glace Bay Beach* (46.177905, -59.9187825) | 1 |
| 2023-05-12 | 1 |
| 2023-06-08 | 0 |
| 2023-07-06 | 0 |
| 2023-07-21 | 0 |
| Cheticamp Beach* (46.600854, -61.03439) | 0 |
| 2023-05-26 | 0 |
| 2023-06-09 | 0 |
| 2023-06-23 | 0 |
| 2023-07-05 | 0 |
| 2023-07-20 | 0 |
| Conrad's Beach (44.64211, -63.36849) | 12 |
| 2023-06-16 | 0 |
| 2023-06-30 | 3 |
| 2023-07-21 | 6 |
| 2023-08-04 | 1 |
| 2023-08-18 | 2 |
| Dominion Beach* (46.21531333, -60.03217667) | 1 |
| 2023-05-12 | 1 |
| 2023-05-24 | 0 |
| 2023-06-08 | 0 |
| 2023-07-06 | 0 |
| 2023-07-21 | 0 |
| Inverness Beach* (46.2319008, -61.3175827) | 3 |
| 2023-05-11 | 1 |
| 2023-05-26 | 0 |
| 2023-06-09 | 0 |
| 2023-06-23 | 2 |
| 2023-07-05 | 0 |
| 2023-07-20 | 0 |
| Lawrencetown Beach (44.64201, -63.32590909) | 16 |
| 2023-06-30 | 7 |
| 2023-07-13 | 4 |
| 2023-07-21 | 4 |
| 2023-08-04 | 0 |
| 2023-08-18 | 1 |
| Margaree Harbour Beach* (46.4440742, -61.1088643) | 3 |
| 2023-05-11 | 2 |
| 2023-05-26 | 0 |
| 2023-06-09 | 0 |
| 2023-06-23 | 1 |
| 2023-07-05 | 0 |
| 2023-07-20 | 0 |
| Martinique Beach (44.68748, -63.10448) | 19 |
| 2023-06-16 | 5 |
| 2023-06-30 | 4 |
| 2023-07-13 | 2 |
| 2023-07-21 | 7 |
| 2023-08-04 | 1 |
| Sable Island- North Beach*(43.9369, -60.0622) | 174 |
| 2023-04-28 | 60 |
| 2023-05-14 | 17 |
| 2023-07-20 | 37 |
| 2023-07-31 | 3 |
| 2023-08-15 | 3 |
| 2023-08-27 | 3 |
| 2023-09-11 |  |
| 2023-09-23 | 1 |
| 2023-10-03 | 50 |
| Sable Island- South Beach* (43.9678, -60.1522) | 302 |
| 2023-04-29 | 26 |
| 2023-05-14 | 2 |
| 2023-05-16 | 25 |
| 2023-07-22 | 149 |
| 2023-08-01 | 31 |
| 2023-08-16 | 17 |
| 2023-08-28 | 8 |
| 2023-09-13 |  |
| 2023-09-24 | 1 |
| 2023-11-03 | 43 |
| **NEWFOUNDLAND** |  |
| Point La Haye* (46.893272, -53.602011) | 34 |
| 2023-05-19 | 27 |
| 2023-06-01 | 1 |
| 2023-06-15 | 3 |
| 2023-06-30 | 0 |
| 2023-07-27 | 3 |
| 2023-08-09 | 0 |
| St. Vincent's* (46.7699613, -53.6130767) | 13 |
| 2023-05-19 | 6 |
| 2023-06-01 | 3 |
| 2023-06-15 | 1 |
| 2023-06-30 | 0 |
| 2023-07-27 | 1 |
| 2023-08-09 | 2 |
| **NEW BRUNSWICK** |  |
| Aboiteau Beach 1 (46.231152, -64.2980815) | 1 |
| 2023-05-08 | 1 |
| 2023-09-08 | 0 |
| Aboiteau Beach 2 (46.2286536, -64.32501845) | 1 |
| 2023-05-08 | 0 |
| 2023-06-22 | 1 |
| Bouctouche Dune* (46.5589481, -63.46193043) | 5 |
| 2023-05-04 | 0 |
| 2023-05-12 | 0 |
| 2023-05-26 | 0 |
| 2023-06-01 | 0 |
| 2023-07-07 | 4 |
| 2023-09-07 | 1 |
| Cap Bimet NB (46.2360135, -64.4566553) | 0 |
| 2023-07-20 | 0 |
| Cap Brule NB (46.237123, -64.4828951) | 0 |
| 2023-07-20 | 0 |
| Cap Lumiere Beach* (46.67568036, -64.71287567) | 9 |
| 2023-05-12 | 0 |
| 2023-06-21 | 2 |
| 2023-07-06 | 4 |
| 2023-08-25 | 3 |
| Escuminac Point (47.07613963, -64.92066663) | 9 |
| 2023-05-26 | 1 |
| 2023-06-22 | 6 |
| 2023-08-03 | 2 |
| Parlee Beach (46.240903, -64.518546) | 1 |
| 2023-05-08 | 1 |
| Petit Chockpish (46.58389315, -64.7202179) | 0 |
| 2023-05-04 | 0 |
| 2023-05-12 | 0 |
| 2023-05-26 | 0 |
| 2023-06-21 | 0 |
| 2023-08-02 | 0 |
| Petit Chockpish (Poteau) (46.60255225, -64.72210105) | 1 |
| 2023-06-01 | 1 |
| 2023-06-21 | 0 |
| Petit-Cap (46.18482965, -64.14825635) | 1 |
| 2023-07-20 | 1 |
| 2023-09-08 | 0 |
| Pointe-Sapin (46.96014725, -64.83632575) | 2 |
| 2023-05-04 | 0 |
| 2023-05-12 | 2 |
| 2023-05-26 | 0 |
| Tracadie Beach* (47.4577841, -64.8770112) | 0 |
| 2023-08-23 | 0 |
| **PRINCE EDWARD ISLAND** |  |
| Black Pond MBS* (46.3612004, -62.1687883) | 0 |
| 2023-06-30 | 0 |
| 2023-07-28 | 0 |
| Cable Head East (46.4665585, -62.5982571) | 0 |
| 2023-07-28 | 0 |
| Cedar Dunes Prov. Park (46.6183096, -64.3848249) | 0 |
| 2023-08-11 | 0 |
| Crowbush Beach* (46.4353693, -62.7998952) | 1 |
| 2023-06-30 | 1 |
| Diligent Pond* (46.4391035, -61.9920746) | 1 |
| 2023-06-29 | 0 |
| 2023-08-29 | 1 |
| Jacques Cartier Park (46.8458806, -64.0162696) | 0 |
| 2023-07-12 | 0 |
| Kildare- Foley's Pond (46.8663029, -63.9978871) | 0 |
| 2023-07-27 | 0 |
| Lower Darnley (46.5621024, -63.6395649) | 2 |
| 2023-07-27 | 2 |
| Nail Pond (46.991772, -64.0742451) | 2 |
| 2023-07-11 | 2 |
| North Lake Harbour (46.4678716, -62.0674166) | 1 |
| 2023-08-30 | 1 |
| Red Point Campground (46.3678621, -62.1382051) | 0 |
| 2023-06-30 | 0 |
| Skinners Pond (46.9548591, -64.1517224) | 3 |
| 2023-07-12 | 3 |
| Stanhope Beach (46.4294295, -63.1385191) | 0 |
| 2023-08-19 | 0 |
| Waterford Campground (46.9538733, -64.1553212) | 0 |
| 2023-07-12 | 0 |

**References**

Chardine, J. W., and G. Pelly. 1994. Operation clean feather: Reducing oil pollution in Newfoundland waters. Technical Report Series, Canadian Wildlife Service, Atlantic Region, Environment Canada. Accessed on February 19, 2024 from <https://publications.gc.ca/collections/collection_2018/eccc/cw69-5/CW69-5-198-eng.pdf>
