## Appendix S3 for "Seabird and sea duck mortalities were lower during the second breeding season in eastern Canada following the introduction of Highly Pathogenic Avian Influenza A H5Nx viruses"

**Appendix S3. A comprehensive breakdown of each group, species, and total reported mortality – across each period**

**Table S1.** Total reported mortality during the fall/winter period (October 1, 2022-March 31, 2023) in five eastern Canadian provinces (New Brunswick, Newfoundland and Labrador, Prince Edward Island, Nova Scotia, and Québec) with assigned species group (waterfowl, seabirds, landbirds, raptors, waders, and shorebirds), common name, number of birds collected for testing, and number of positive HPAI cases detected from testing data collected in November 2023.

| **Species Group** | **Common Name** | **Total Birds** | **% of Period Total** | **Number Tested** | **Number Positive** |
| --- | --- | --- | --- | --- | --- |
| **waterfowl** | Snow Goose | 1643 | 45.72 | 68 | 63 |
| **waterfowl** | Canada Goose | 349 | 9.71 | 87 | 52 |
| **landbirds** | American Crow | 210 | 5.84 | 115 | 40 |
| **seabirds** | Northern Gannet | 206 | 5.73 | 8 | 4 |
| **raptors** | Barred Owl | 126 | 3.51 | 37 | 0 |
| **seabirds** | Herring Gull | 86 | 2.39 | 28 | 6 |
| **seabirds** | Unknown Gull | 55 | 1.53 | 3 | 1 |
| **seabirds** | Thick-billed Murre | 52 | 1.45 | 11 | 0 |
| **raptors** | Northern Saw-whet Owl | 49 | 1.36 | 9 | 0 |
| **waterfowl** | Mallard | 42 | 1.17 | 12 | 4 |
| **landbirds** | Ruffed Grouse | 42 | 1.17 | 6 | 0 |
| **landbirds** | European Starling | 40 | 1.11 | 6 | 0 |
| **raptors** | Bald Eagle | 39 | 1.09 | 16 | 6 |
| **waterfowl** | American Black Duck | 34 | 0.95 | 12 | 12 |
| **seabirds** | Great Black-backed Gull | 31 | 0.86 | 11 | 2 |
| **seabirds** | Northern Fulmar | 25 | 0.7 | 0 | 0 |
| **raptors** | Red-tailed Hawk | 25 | 0.7 | 17 | 6 |
| **landbirds** | Rock Pigeon | 24 | 0.67 | 3 | 1 |
| **raptors** | Great Horned Owl | 19 | 0.53 | 8 | 1 |
| **raptors** | Cooper's Hawk | 18 | 0.5 | 12 | 0 |
| **seabirds** | Iceland Gull | 18 | 0.5 | 2 | 1 |
| **seabirds** | Black-legged Kittiwake | 17 | 0.47 | 2 | 1 |
| **seabirds** | Dovekie | 17 | 0.47 | 3 | 0 |
| **seabirds** | Atlantic Puffin | 15 | 0.42 | 4 | 0 |
| **seabirds** | Double-crested Cormorant | 15 | 0.42 | 12 | 0 |
| **landbirds** | Black-capped Chickadee | 14 | 0.39 | 0 | 0 |
| **loons** | Common Loon | 14 | 0.39 | 4 | 0 |
| **seabirds** | Ring-billed Gull | 14 | 0.39 | 9 | 2 |
| **landbirds** | Blue Jay | 13 | 0.36 | 6 | 0 |
| **landbirds** | Ring-necked Pheasant | 13 | 0.36 | 0 | 0 |
| **raptors** | Sharp-shinned Hawk | 13 | 0.36 | 6 | 0 |
| **landbirds** | American Goldfinch | 12 | 0.33 | 3 | 0 |
| **seabirds** | Cory's Shearwater | 12 | 0.33 | 12 | 0 |
| **raptors** | Merlin | 11 | 0.31 | 9 | 0 |
| **raptors** | Peregrine Falcon | 11 | 0.31 | 7 | 4 |
| **seabirds** | Common Murre | 10 | 0.28 | 2 | 0 |
| **raptors** | Unknown Hawk | 10 | 0.28 | 0 | 0 |
| **landbirds** | Wild Turkey | 9 | 0.25 | 6 | 0 |
| **waterfowl** | Common Goldeneye | 8 | 0.22 | 2 | 1 |
| **landbirds** | Mourning Dove | 8 | 0.22 | 1 | 0 |
| **raptors** | Turkey Vulture | 8 | 0.22 | 5 | 1 |
| **raptors** | Unknown Owl | 8 | 0.22 | 0 | 0 |
| **seabirds** | Unknown Alcid (Alcidae sp.) | 7 | 0.19 | 0 | 0 |
| **seabirds** | Common Eider | 6 | 0.17 | 5 | 3 |
| **waterfowl** | Common Merganser | 6 | 0.17 | 0 | 0 |
| **landbirds** | Dark-eyed Junco | 7 | 0.17 | 0 | 0 |
| **seabirds** | Great Shearwater | 6 | 0.17 | 6 | 0 |
| **waterfowl** | Surf Scoter | 6 | 0.17 | 4 | 0 |
| **raptors** | American Kestrel | 5 | 0.14 | 1 | 0 |
| **raptors** | Boreal Owl | 5 | 0.14 | 0 | 0 |
| **landbirds** | Common Grackle | 5 | 0.14 | 0 | 0 |
| **landbirds** | Common Raven | 5 | 0.14 | 3 | 2 |
| **raptors** | Golden Eagle | 5 | 0.14 | 5 | 0 |
| **waders** | Great Blue Heron | 5 | 0.14 | 3 | 0 |
| **seabirds** | Manx Shearwater | 5 | 0.14 | 5 | 0 |
| **landbirds** | Northern Cardinal | 5 | 0.14 | 1 | 1 |
| **waterfowl** | Unknown Duck | 5 | 0.14 | 1 | 0 |
| **landbirds** | American Robin | 4 | 0.11 | 0 | 0 |
| **waterfowl** | Black Scoter | 4 | 0.11 | 1 | 0 |
| **landbirds** | Bohemian Waxwing | 4 | 0.11 | 1 | 0 |
| **landbirds** | Cedar Waxwing | 4 | 0.11 | 0 | 0 |
| **seabirds** | Common Tern | 4 | 0.11 | 4 | 0 |
| **landbirds** | Golden-crowned Kinglet | 4 | 0.11 | 0 | 0 |
| **landbirds** | House Finch | 4 | 0.11 | 0 | 0 |
| **landbirds** | House Sparrow | 4 | 0.11 | 2 | 1 |
| **landbirds** | Pine Grosbeak | 4 | 0.11 | 0 | 0 |
| **seabirds** | Sooty Shearwater | 4 | 0.11 | 3 | 0 |
| **landbirds** | Evening Grosbeak | 3 | 0.08 | 0 | 0 |
| **landbirds** | Gray Partridge | 3 | 0.08 | 0 | 0 |
| **landbirds** | Hairy Woodpecker | 3 | 0.08 | 0 | 0 |
| **waterfowl** | Red-breasted Merganser | 3 | 0.08 | 0 | 0 |
| **raptors** | Snowy Owl | 3 | 0.08 | 3 | 0 |
| **landbirds** | Unknown Sparrow | 3 | 0.08 | 0 | 0 |
| **landbirds** | White-throated Sparrow | 3 | 0.08 | 0 | 0 |
| **waterfowl** | Wood Duck | 3 | 0.08 | 2 | 0 |
| **shorebirds** | American Woodcock | 2 | 0.06 | 0 | 0 |
| **waterfowl** | Blue-winged Teal | 2 | 0.06 | 0 | 0 |
| **raptors** | Broad-winged Hawk | 2 | 0.06 | 1 | 0 |
| **waterfowl** | Cackling Goose | 2 | 0.06 | 2 | 2 |
| **landbirds** | Common Redpoll | 2 | 0.06 | 0 | 0 |
| **raptors** | Eastern Screech-Owl | 2 | 0.06 | 0 | 0 |
| **seabirds** | Glaucous Gull | 2 | 0.06 | 1 | 0 |
| **seabirds** | Great Cormorant | 2 | 0.06 | 0 | 0 |
| **waterfowl** | Green-winged Teal | 2 | 0.06 | 0 | 0 |
| **landbirds** | Lincoln's Sparrow | 2 | 0.06 | 0 | 0 |
| **raptors** | Northern Goshawk | 2 | 0.06 | 2 | 0 |
| **raptors** | Osprey | 2 | 0.06 | 2 | 0 |
| **waders** | Purple Gallinule | 2 | 0.06 | 0 | 0 |
| **waterfowl** | Ross's Goose | 2 | 0.06 | 0 | 0 |
| **raptors** | Rough-legged Hawk | 2 | 0.06 | 1 | 0 |
| **shorebirds** | Semipalmated Sandpiper | 2 | 0.06 | 1 | 0 |
| **landbirds** | Song Sparrow | 2 | 0.06 | 0 | 0 |
| **Unknown Bird** | Unknown Bird | 2 | 0.06 | 0 | 0 |
| **landbirds** | White-breasted Nuthatch | 2 | 0.06 | 0 | 0 |
| **shorebirds** | White-rumped Sandpiper | 2 | 0.06 | 0 | 0 |
| **waders** | American Coot | 1 | 0.03 |  |  |
| **landbirds** | American Pipit | 1 | 0.03 | 0 | 0 |
| **landbirds** | Bank Swallow | 1 | 0.03 | 1 | 0 |
| **landbirds** | Blue-gray Gnatcatcher | 1 | 0.03 | 0 | 0 |
| **waterfowl** | Brant | 1 | 0.03 | 0 | 0 |
| **landbirds** | Chipping Sparrow | 1 | 0.03 | 0 | 0 |
| **landbirds** | Downy Woodpecker | 1 | 0.03 | 0 | 0 |
| **waterfowl** | Greater Scaup | 1 | 0.03 | 0 | 0 |
| **landbirds** | Hermit Thrush | 1 | 0.03 | 0 | 0 |
| **waterfowl** | Hooded Merganser | 1 | 0.03 | 1 | 0 |
| **seabirds** | Leach's Storm-Petrel | 1 | 0.03 | 1 | 1 |
| **waterfowl** | Mute Swan | 1 | 0.03 | 1 | 1 |
| **landbirds** | Northern Flicker | 1 | 0.03 | 0 | 0 |
| **landbirds** | Northern Mockingbird | 1 | 0.03 | 1 | 0 |
| **landbirds** | Orange-crowned Warbler | 1 | 0.03 | 0 | 0 |
| **landbirds** | Red-breasted Nuthatch | 1 | 0.03 | 0 | 0 |
| **waders** | Sandhill Crane | 1 | 0.03 | 0 | 0 |
| **raptors** | Short-eared Owl | 1 | 0.03 | 1 | 0 |
| **landbirds** | Spruce Grouse | 1 | 0.03 | 0 | 0 |
| **seabirds** | Unknown Cormorant | 1 | 0.03 | 0 | 0 |
| **landbirds** | Unknown Woodpecker | 1 | 0.03 | 0 | 0 |
| **landbirds** | Yellow-billed Cuckoo | 1 | 0.03 | 0 | 0 |

**Table S2.** Total reported mortality during the spring/summer period (April 1, 2023-September 30, 2023) in five eastern Canadian provinces (New Brunswick, Newfoundland and Labrador, Prince Edward Island, Nova Scotia, and Québec) with assigned species group (waterfowl, seabirds, landbirds, raptors, waders, and shorebirds), common name, number of birds collected for testing, and number of positive HPAI cases detected from testing data collected in November 2023.

| **Species Group** | **Common Name** | **Total Birds** | **% of Period Total** | **Number Tested** | **Number Positive** |
| --- | --- | --- | --- | --- | --- |
| **landbirds** | American Crow | 298 | 8.81 | 98 | 11 |
| **seabirds** | Great Shearwater | 282 | 8.34 | 7 | 0 |
| **seabirds** | Northern Gannet | 156 | 4.61 | 25 | 4 |
| **seabirds** | Herring Gull | 153 | 4.52 | 32 | 5 |
| **seabirds** | Unknown Gull | 152 | 4.49 | 1 | 0 |
| **waterfowl** | Canada Goose | 148 | 4.37 | 51 | 3 |
| **seabirds** | Common Murre | 145 | 4.29 | 4 | 0 |
| **seabirds** | Leach's Storm-Petrel | 114 | 3.37 | 24 | 0 |
| **raptors** | Bald Eagle | 105 | 3.1 | 53 | 5 |
| **seabirds** | Great Black-backed Gull | 81 | 2.39 | 16 | 7 |
| **seabirds** | Common Tern | 72 | 2.13 | 56 | 1 |
| **waders** | Great Blue Heron | 64 | 1.89 | 17 | 0 |
| **seabirds** | Thick-billed Murre | 64 | 1.89 | 16 | 0 |
| **landbirds** | American Robin | 63 | 1.86 | 0 | 0 |
| **waterfowl** | Snow Goose | 58 | 1.71 | 10 | 1 |
| **landbirds** | Rock Pigeon | 49 | 1.45 | 8 | 0 |
| **waterfowl** | Mallard | 48 | 1.42 | 16 | 0 |
| **raptors** | Merlin | 43 | 1.27 | 2 | 0 |
| **landbirds** | European Starling | 42 | 1.24 | 2 | 0 |
| **seabirds** | Double-crested Cormorant | 39 | 1.15 | 12 | 0 |
| **seabirds** | Northern Fulmar | 37 | 1.09 | 13 | 12 |
| **seabirds** | Ring-billed Gull | 37 | 1.09 | 17 | 1 |
| **seabirds** | Common Eider | 35 | 1.03 | 12 | 0 |
| **seabirds** | Unknown Tern | 35 | 1.03 | 13 | 0 |
| **raptors** | Barred Owl | 34 | 1.01 | 5 | 1 |
| **seabirds** | Unknown Alcid (Alcidae sp.) | 34 | 1.01 | 0 | 0 |
| **waterfowl** | Unknown Duck | 34 | 1.01 | 5 | 0 |
| **landbirds** | Blue Jay | 31 | 0.92 | 4 | 1 |
| **landbirds** | Common Grackle | 30 | 0.89 | 0 | 0 |
| **raptors** | Red-tailed Hawk | 29 | 0.86 | 18 | 3 |
| **landbirds** | Common Raven | 28 | 0.83 | 6 | 0 |
| **loons** | Common Loon | 25 | 0.74 | 8 | 0 |
| **landbirds** | Wild Turkey | 23 | 0.68 | 8 | 0 |
| **loons** | Red-throated Loon | 22 | 0.65 | 7 | 0 |
| **landbirds** | Ruffed Grouse | 21 | 0.62 | 2 | 0 |
| **seabirds** | Sooty Shearwater | 21 | 0.62 | 2 | 0 |
| **seabirds** | Black-legged Kittiwake | 20 | 0.59 | 4 | 1 |
| **raptors** | Great Horned Owl | 19 | 0.56 | 8 | 0 |
| **seabirds** | Unknown Cormorant | 19 | 0.56 | 0 | 0 |
| **raptors** | Osprey | 18 | 0.53 | 4 | 0 |
| **raptors** | Peregrine Falcon | 18 | 0.53 | 5 | 0 |
| **raptors** | Unknown Owl | 18 | 0.53 | 0 | 0 |
| **raptors** | Turkey Vulture | 16 | 0.47 | 9 | 3 |
| **Unknown Bird** | Unknown Bird | 16 | 0.47 | 0 | 0 |
| **landbirds** | Unknown Sparrow | 15 | 0.44 | 0 | 0 |
| **landbirds** | American Goldfinch | 14 | 0.41 | 0 | 0 |
| **seabirds** | Atlantic Puffin | 14 | 0.41 | 6 | 0 |
| **raptors** | Red-shouldered Hawk | 14 | 0.41 | 0 | 0 |
| **raptors** | Sharp-shinned Hawk | 14 | 0.41 | 2 | 0 |
| **raptors** | American Kestrel | 13 | 0.38 | 1 | 0 |
| **seabirds** | Roseate Tern | 13 | 0.38 | 5 | 0 |
| **landbirds** | Unknown Woodpecker | 13 | 0.38 | 0 | 0 |
| **seabirds** | Black Guillemot | 12 | 0.35 | 3 | 0 |
| **seabirds** | Dovekie | 12 | 0.35 | 2 | 0 |
| **landbirds** | Ruby-throated Hummingbird | 12 | 0.35 | 2 | 0 |
| **landbirds** | Dark-eyed Junco | 11 | 0.33 | 0 | 0 |
| **raptors** | Northern Saw-whet Owl | 11 | 0.33 | 0 | 0 |
| **landbirds** | Black-capped Chickadee | 9 | 0.27 | 0 | 0 |
| **raptors** | Broad-winged Hawk | 9 | 0.27 | 0 | 0 |
| **raptors** | Cooper's Hawk | 9 | 0.27 | 0 | 0 |
| **seabirds** | Iceland Gull | 9 | 0.27 | 0 | 0 |
| **landbirds** | Northern Flicker | 9 | 0.27 | 0 | 0 |
| **raptors** | Northern Goshawk | 9 | 0.27 | 1 | 0 |
| **loons** | Unknown Loon | 9 | 0.27 | 0 | 0 |
| **shorebirds** | American Woodcock | 8 | 0.24 | 0 | 0 |
| **landbirds** | Mourning Dove | 8 | 0.24 | 0 | 0 |
| **landbirds** | Ovenbird | 8 | 0.24 | 0 | 0 |
| **seabirds** | Razorbill | 8 | 0.24 | 0 | 0 |
| **seabirds** | Unknown Murre | 8 | 0.24 | 0 | 0 |
| **seabirds** | Arctic Tern | 7 | 0.21 | 7 | 0 |
| **landbirds** | Cedar Waxwing | 7 | 0.21 | 0 | 0 |
| **landbirds** | Ring-necked Pheasant | 7 | 0.21 | 1 | 0 |
| **landbirds** | Song Sparrow | 7 | 0.21 | 0 | 0 |
| **raptors** | Unknown Hawk | 7 | 0.21 | 0 | 0 |
| **landbirds** | Black-and-white Warbler | 6 | 0.18 | 0 | 0 |
| **raptors** | Chinese Sparrowhawk | 6 | 0.18 | 0 | 0 |
| **landbirds** | Gray Partridge | 6 | 0.18 | 0 | 0 |
| **seabirds** | Manx Shearwater | 6 | 0.18 | 3 | 0 |
| **landbirds** | Pileated Woodpecker | 6 | 0.18 | 0 | 0 |
| **landbirds** | Red-eyed Vireo | 6 | 0.18 | 0 | 0 |
| **waterfowl** | Wood Duck | 6 | 0.18 | 4 | 0 |
| **waterfowl** | American Black Duck | 5 | 0.15 | 0 | 0 |
| **landbirds** | Barn Swallow | 5 | 0.15 | 0 | 0 |
| **raptors** | Boreal Owl | 5 | 0.15 | 0 | 0 |
| **landbirds** | Chimney Swift | 5 | 0.15 | 1 | 0 |
| **landbirds** | Common Yellowthroat | 5 | 0.15 | 0 | 0 |
| **landbirds** | Downy Woodpecker | 5 | 0.15 | 0 | 0 |
| **shorebirds** | Killdeer | 5 | 0.15 | 1 | 0 |
| **landbirds** | Purple Finch | 5 | 0.15 | 0 | 0 |
| **Unknown** | Unknown Bird | 5 | 0.15 | 0 | 0 |
| **landbirds** | White-throated Sparrow | 5 | 0.15 | 0 | 0 |
| **landbirds** | American Redstart | 4 | 0.12 | 0 | 0 |
| **landbirds** | Belted Kingfisher | 4 | 0.12 | 1 | 0 |
| **waterfowl** | Common Goldeneye | 4 | 0.12 | 1 | 0 |
| **landbirds** | Common Nighthawk | 4 | 0.12 | 2 | 0 |
| **landbirds** | Northern Parula | 4 | 0.12 | 0 | 0 |
| **raptors** | Snowy Owl | 4 | 0.12 | 3 | 1 |
| **shorebirds** | Spotted Sandpiper | 4 | 0.12 | 0 | 0 |
| **waterfowl** | Surf Scoter | 4 | 0.12 | 1 | 0 |
| **landbirds** | Swainson's Thrush | 4 | 0.12 | 0 | 0 |
| **seabirds** | Unknown Shearwater | 4 | 0.12 | 0 | 0 |
| **landbirds** | Unknown Swallow | 4 | 0.12 | 0 | 0 |
| **landbirds** | Yellow-bellied Sapsucker | 4 | 0.12 | 0 | 0 |
| **waders** | American Bittern | 3 | 0.09 | 0 | 0 |
| **landbirds** | American Pipit | 3 | 0.09 | 0 | 0 |
| **landbirds** | Golden-crowned Kinglet | 3 | 0.09 | 0 | 0 |
| **landbirds** | Gray Catbird | 3 | 0.09 | 1 | 0 |
| **landbirds** | Nashville Warbler | 3 | 0.09 | 0 | 0 |
| **raptors** | Northern Harrier | 3 | 0.09 | 0 | 0 |
| **landbirds** | Pine Grosbeak | 3 | 0.09 | 0 | 0 |
| **landbirds** | Red-winged Blackbird | 3 | 0.09 | 0 | 0 |
| **landbirds** | Song Thrush | 3 | 0.09 | 0 | 0 |
| **landbirds** | Tennessee Warbler | 3 | 0.09 | 0 | 0 |
| **waterfowl** | Trumpeter Swan | 3 | 0.09 | 1 | 1 |
| **landbirds** | Unknown Warbler | 3 | 0.09 | 0 | 0 |
| **landbirds** | Veery | 3 | 0.09 | 0 | 0 |
| **landbirds** | Winter Wren | 3 | 0.09 | 0 | 0 |
| **landbirds** | Yellow Warbler | 3 | 0.09 | 0 | 0 |
| **waterfowl** | American Wigeon | 2 | 0.06 | 0 | 0 |
| **landbirds** | Blackpoll Warbler | 2 | 0.06 | 0 | 0 |
| **landbirds** | Black-throated Blue Warbler | 2 | 0.06 | 0 | 0 |
| **waterfowl** | Blue-winged Teal | 2 | 0.06 | 1 | 0 |
| **landbirds** | Bohemian Waxwing | 2 | 0.06 | 1 | 0 |
| **landbirds** | Cape May Warbler | 2 | 0.06 | 0 | 0 |
| **landbirds** | Chestnut-sided Warbler | 2 | 0.06 | 0 | 0 |
| **waterfowl** | Common Merganser | 2 | 0.06 | 0 | 0 |
| **landbirds** | Common Redpoll | 2 | 0.06 | 0 | 0 |
| **seabirds** | Cory's Shearwater | 2 | 0.06 | 0 | 0 |
| **landbirds** | Eastern Phoebe | 2 | 0.06 | 0 | 0 |
| **landbirds** | Fox Sparrow | 2 | 0.06 | 0 | 0 |
| **waterfowl** | Gadwall | 2 | 0.06 | 1 | 0 |
| **seabirds** | Great Cormorant | 2 | 0.06 | 0 | 0 |
| **waterfowl** | Greater Scaup | 2 | 0.06 | 0 | 0 |
| **landbirds** | House Sparrow | 2 | 0.06 | 0 | 0 |
| **landbirds** | Magnolia Warbler | 2 | 0.06 | 0 | 0 |
| **landbirds** | Northern Cardinal | 2 | 0.06 | 0 | 0 |
| **landbirds** | Red Crossbill | 2 | 0.06 | 0 | 0 |
| **landbirds** | Rose-breasted Grosbeak | 2 | 0.06 | 0 | 0 |
| **raptors** | Rough-legged Hawk | 2 | 0.06 | 0 | 0 |
| **shorebirds** | Semipalmated Plover | 2 | 0.06 | 0 | 0 |
| **landbirds** | Tree Swallow | 2 | 0.06 | 0 | 0 |
| **landbirds** | Tufted Titmouse | 2 | 0.06 | 0 | 0 |
| **landbirds** | American Goshawk | 1 | 0.03 | 0 | 0 |
| **shorebirds** | Baird's Sandpiper | 1 | 0.03 | 0 | 0 |
| **waterfowl** | Black Brant | 1 | 0.03 | 1 | 0 |
| **waterfowl** | Black Scoter | 1 | 0.03 | 0 | 0 |
| **shorebirds** | Black-bellied Plover | 1 | 0.03 | 0 | 0 |
| **landbirds** | Black-billed Cuckoo | 1 | 0.03 | 0 | 0 |
| **landbirds** | Blackburnian Warbler | 1 | 0.03 | 0 | 0 |
| **waders** | Black-crowned Night-Heron | 1 | 0.03 | 0 | 0 |
| **landbirds** | Black-throated Green Warbler | 1 | 0.03 | 0 | 0 |
| **seabirds** | Brown Booby | 1 | 0.03 | 0 | 0 |
| **landbirds** | Brown-headed Cowbird | 1 | 0.03 | 1 | 0 |
| **waterfowl** | Cackling Goose | 1 | 0.03 | 1 | 0 |
| **landbirds** | Canada Warbler | 1 | 0.03 | 0 | 0 |
| **landbirds** | Chipping Sparrow | 1 | 0.03 | 0 | 0 |
| **landbirds** | Cliff Swallow | 1 | 0.03 | 0 | 0 |
| **waders** | Common Gallinule | 1 | 0.03 | 0 | 0 |
| **landbirds** | Eastern Bluebird | 1 | 0.03 | 0 | 0 |
| **landbirds** | Eastern Kingbird | 1 | 0.03 | 0 | 0 |
| **landbirds** | Eastern Whip-poor-will | 1 | 0.03 | 0 | 0 |
| **seabirds** | Glaucous Gull | 1 | 0.03 | 0 | 0 |
| **raptors** | Golden Eagle | 1 | 0.03 | 0 | 0 |
| **landbirds** | Ipswich Sparrow | 1 | 0.03 | 0 | 0 |
| **seabirds** | Lesser Black-backed Gull | 1 | 0.03 | 0 | 0 |
| **shorebirds** | Lesser Yellowlegs | 1 | 0.03 | 0 | 0 |
| **landbirds** | Mourning Warbler | 1 | 0.03 | 0 | 0 |
| **landbirds** | Northern Mockingbird | 1 | 0.03 | 0 | 0 |
| **landbirds** | Northern Shrike | 1 | 0.03 | 0 | 0 |
| **landbirds** | Northern Waterthrush | 1 | 0.03 | 0 | 0 |
| **landbirds** | Pine Warbler | 1 | 0.03 | 0 | 0 |
| **seabirds** | Pomarine Jaeger | 1 | 0.03 | 0 | 0 |
| **landbirds** | Purple Martin | 1 | 0.03 | 0 | 0 |
| **waterfowl** | Red-breasted Merganser | 1 | 0.03 | 0 | 0 |
| **shorebirds** | Ruddy Turnstone | 1 | 0.03 | 0 | 0 |
| **landbirds** | Savannah Sparrow | 1 | 0.03 | 0 | 0 |
| **raptors** | Short-eared Owl | 1 | 0.03 | 0 | 0 |
| **waders** | Sora | 1 | 0.03 | 0 | 0 |
| **landbirds** | Spruce Grouse | 1 | 0.03 | 0 | 0 |
| **landbirds** | Swamp Sparrow | 1 | 0.03 | 0 | 0 |
| **landbirds** | Unknown Passerine | 1 | 0.03 | 0 | 0 |
| **landbirds** | Unknown Thrush | 1 | 0.03 | 0 | 0 |
| **waders** | Virginia Rail | 1 | 0.03 | 0 | 0 |
| **landbirds** | Warbling Vireo | 1 | 0.03 | 0 | 0 |
| **landbirds** | White-breasted Nuthatch | 1 | 0.03 | 0 | 0 |
| **shorebirds** | Wilson's Snipe | 1 | 0.03 | 0 | 0 |
| **seabirds** | Wilson's Storm-Petrel | 1 | 0.03 | 0 | 0 |
| **landbirds** | Yellow-bellied Flycatcher | 1 | 0.03 | 0 | 0 |
| **landbirds** | Yellow-throated Vireo | 1 | 0.03 | 0 | 0 |
