## Appendix S4 for "Seabird and sea duck mortalities were lower during the second breeding season in eastern Canada following the introduction of Highly Pathogenic Avian Influenza A H5Nx viruses"

**Appendix S4. Additional Acknowledgements**

Most reported mortalities can be traced back to engaged members of the public, and we offer our sincere thanks to all individuals who shared their observations. Numerous individuals from Indigenous, federal, provincial, and municipal governments as well as non-governmental organizations have been instrumental in the completion of this research. Thank you.

**Indigenous partners:**
This research was conducted in what is now known as eastern Canada on the traditional land of the Mi’kmaq First Nation, the Wolastoqiyik (Maliseet) First Nation, the Passamaquoddy people, the Beothuk peoples, the Innu of Nitassinan, the Inuit of Nunatsiavut, and the southern Inuit of NunatuKavut. We would like to acknowledge the support provided by the staff and members of the Innu Nation, the Miawpukek First Nation (MFN), the Nunatsiavut Government, the NunatuKavut Community Council, and the Qalipu First Nation.

**Government Agencies and NGOs:**

We would like to acknowledge the invaluable support provided by staff from the following organizations:

Canadian Wildlife Health Cooperative (CWHC- Atlantic, CWHC - Québec),

Canadian Food Inspection Agency, National Centre of Foreign Animal Disease, Environment and Climate Change Canada, Newfoundland and Labrador Department of Fisheries, Forestry and Agriculture (including Regional Services, Animal Health Division, Natural Areas Division and Wildlife Division), Nova Scotia Department of Natural Resources and Renewables (Wildlife Division), Prince Edward Island Department of Environment, Water and Climate Change, Forests, Fish and Wildlife Division, Municipality of the Magdalen Islands (Les Iles-de-la-Madeleine), Ministère de l’Environnement, de la Lutte contre les changements climatiques, de la Faune et des Parcs du Québec, Ministère de l'Agriculture, des Pêcheries et de l'Alimentation du Québec, Birds Canada, Nature NB, Canadian Parks and Wilderness Society - The Newfoundland and Labrador Chapter (CPAWS - NL), Indian Bay Ecosystems Initiative, Rock Wildlife Rescue, Société Duvetnor Ltée, Société protectrice des eiders de l’estuaire, Société Provancher, Parc National de l’Île-Bonaventure-et-du-Rocher-Percé, Parks Canada (including Mingan Archipelago National Park Reserve, Prince Edward Island National Park, Kouchibouguac National Park, Cape Breton Highlands National Park, Saguenay-St. Lawrence Marine Park, Kejimkujik National Park, Forillon National Park), Fisheries and Oceans Canada, Transport Canada’s National Aerial Surveillance Program, and Sable Island Institute.

**Individuals:**

Our deepest appreciation goes to the following individuals who contributed mortality data to this study:

Kathleen Aikens, Meghan Baker, Audrée Benoit, Matthieu Beaumont, Yohannes Berhanes, Katherine Bond, Michael Brown, Kevin Craig, Shawn Craik, Gail Davoren, Gabrielle Dimitri-Masson, Thibaud Durbecq, Suzanne Dooley, Alexis Godin, Daniel Gallant, Karen Gosse, Carina Gjerdrum, Jean-François Giroux, Garry Gregory, Megan Fortune, Drew Hutchinson, Gregory Jeddore, Anabelle Hebert, Ted Leighton, François Lapointe, Raphaël A. Lavoie, Zoe Lucas, Heather Major, Dave McRuer, Mark Mallory, Mark Maddox, Chris Mooney, René Nault, Ingrid Pollet, Sydney Collins, Jennifer Provencher, Jean-Simon Richard, Lewnanny Richardson, Anna Robuck, Jennifer Rock, Frank Shapleigh, Christopher Poole, Sydney Collins, Glen Parsons, Chris Ward, Regina Wells, Christine Lepage, Liam Taylor, Georgia Taylor, Zoe Lucas, Andrew Kennedy, Suzanne Dooley, Ted Leighton, Christopher Poole, Trevor Thompson, Anna Robuck.
